# Antagonizing niche signals regulate the emergence of postnatal neural stem cells and allow enlargement of their pool

**DOI:** 10.64898/2026.07.09.737275

**Authors:** D. Cimino, A. Danese, A. Denoth-Lippuner, P. Chlebik, M. Thorwirth, T. Simon, M. Richter, S. Jessberger, M. Götz

## Abstract

Adult neural stem cells (aNSCs) of the lateral ventricular sub-ventricular zone (V-SVZ) are set aside during embryogenesis from a pool of slowly dividing neural stem/progenitor cells (NSPCs) residing in the lateral ganglionic eminence (LGE). The time of aNSCs specification coincides with the peak of embryonic neurogenesis, raising the question of which mechanisms allow only a subpopulation of NSPCs to retain stem cell identity while others proceed to divide and differentiate into neurons. To address this, we isolated rapidly and slowly dividing NSPCs from the mouse LGE at mid-neurogenesis, when aNSCs are specified, and profiled the transcriptome at the single cell level. We find that slowly dividing NSPCs constitute a heterogeneous population encompassing both radial glia cells (RGCs) and intermediate progenitors (IPs) and characterized by different gene signatures. Focusing on RGCs, we show that slowly and rapidly dividing RGCs differentially express genes involved in the formation – such as *Lamb2* – or degradation – such as *Mmp15* – of the extracellular matrix (ECM), pointing to opposite niche-remodelling strategies as a potential mechanism of fate determination. *In vivo* perturbation of *Lamb2* and *Mmp15* affects the seeding and maintenance of NSCs in an opposing manner, with knockdown of *Mmp15* increasing the NSC pool as also shown by single-cell RNA-sequencing. These findings suggest that opposite ECM remodelling by slowly and rapidly dividing RGCs constitutes a novel cross-regulatory niche-level mechanism regulating aNSCs emergence, opening new avenues for regulating their numbers and thereby long-term maintenance of the brain’s regenerative potential.

## Introduction

In most regions of the mammalian brain, neurogenesis occurs predominantly during embryogenesis and terminates around the time of birth. However, in some areas of the adult brain, new neurons continue to be produced by aNSCs residing in restricted and specialized niches (Doetsch et al., 1999; Kempermann et al., 2003). In mice, the largest of the adult neurogenic niches is the V-SVZ, corresponding to the most apical periventricular area of the lateral ventricles. Here, aNSCs sustain neurogenesis for the olfactory bulb (OB) through the production of different interneuron subtypes (Fuentealba et al., 2015; Merkle et al., 2014). In addition, aNSCs generate glial cells and the gliogenic/neurogenic potential of aNSCs differs among the dorsal, medial and lateral walls of the V-SVZ, with the lateral wall being the most neurogenic domain (Ortega et al., 2013; Mizrak et al., 2019).

The majority of the aNSCs of the lateral V-SVZ emerges early in development, during a timeframe coinciding with the peak of embryonic neurogenesis (Fuentealba et al., 2015; Furutachi et al., 2015). Labelling-retention assays have indeed demonstrated that aNSCs of the lateral V-SVZ originate from a subset of NSPCs, residing in the LGE, which slow down their cell-cycle around the embryonic days (E) 13-16 to eventually become quiescent until postnatal life (Fuentealba et al., 2015; Furutachi et al., 2015). This raises the intriguing question of how slowly dividing aNSCs ancestors retain their stem cell identity throughout development, while neurogenic NSPCs located in the same niche continue dividing and differentiating to drive embryonic neurogenesis.

In the adult V-SVZ, the balance between NSCs quiescence and activation is tightly regulated by the surrounding environment. Adult NSCs are embedded in a complex niche where blood vessels, growth factors and ECM components form an integrated regulatory system that simultaneously drives the proliferation and differentiation of activated NSCs while preserving the quiescence of others (Basak et al., 2012; Douet et al., 2012; Kazanis et al., 2010; Lim et al., 2000a; Nascimento et al., 2018; Porlan et al., 2014; Shan et al., 2018; Tavazoie et al., 2008). Similarly, in the embryonic brain, different cues from the surrounding microenvironment might drive the differentiation of rapidly dividing NSPCs, while promoting the maintenance of an undifferentiated state in slowly dividing NSPCs, thereby allowing the establishment of the aNSCs pool and the continuation of embryonic neurogenesis. To date, the mechanisms underlying the emergence of aNSCs in development are not fully understood, and the role of the extracellular environment in this process is largely unexplored.

To address this gap, we transcriptomically profiled slowly and rapidly dividing LGE NSPCs at mid-neurogenesis, uncovering differential expression of ECM-associated genes between these populations and a novel niche-mediated regulation of aNSC specification.

## Results

### Isolation and transcriptomic profiling of slowly and rapidly dividing LGE NSPCs

To investigate the mechanisms underlying aNSCs emergence, we profiled the transcriptome of slowly and rapidly dividing NSPCs from the mouse LGE – the embryonic region generating most of the lateral V-SVZ aNSCs – at E16, i.e. the close of the time window of aNSCs specification (Fuentealba et al., 2015; Furutachi et al., 2015). To do so, we used the iCOUNT transgenic mice, in which the cell proliferation rate can be inferred by retention of H3.1-mCherry and H3.1-GFP fusion proteins (Denoth-Lippuner et al., 2021). Briefly, in the iCOUNT mice, *mCherry* and *GFP* are fused to the C-terminus of the cell-cycle dependent histone variant *H3.1*. The *mCherry* coding sequence and a downstream STOP codon are flanked by loxP sites so that, in the absence of Cre recombinase, the H3.1-mCherry fusion protein is expressed at each cell division, while upon Cre-mediated recombination of the loxP sites, expression is switched to the GFP-tagged H3.1, replacing the mCherry-tagged H3.1 in a cell division-dependant manner. (Fig. 1A). To temporally control the tracking of the cell division rate, iCOUNT mice were crossed with a Rosa26Cre-ERT2 mouse line, carrying the Cre recombinase fused to the estrogen receptor T2, which prevents the activation of the recombinase in absence of tamoxifen and expression of the Cre-ERT2 fusion protein is driven by the ubiquitous promoter Rosa26 (Ventura et al., 2007). Excision of the floxed *mCherry* sequence, and subsequent expression of the GFP-tagged histone, was induced at E14 by tamoxifen injection. Animals were sacrificed two days post injection to allow dilution of the H3.1-mCherry fusion protein in highly proliferative cells (Fig. 1B). Slowly (mCherry+ GFP+) and rapidly (GFP+) dividing cells were isolated via fluorescent-activated cell sorting (FACS) (Fig. 1B). To exclude possible non-recombined cells, single mCherry+ cells were discarded. Cells were collected into 384-well plates and sequenced using VASA-seq technology (Fig. 1C), a single-cell full-length RNA-sequencing method that shows higher sensitivity and gene detection rate as compared to other RNA-sequencing methods, thereby improving the detection of lowly expressed transcripts (Salmen et al., 2022).

**Figure 1.**
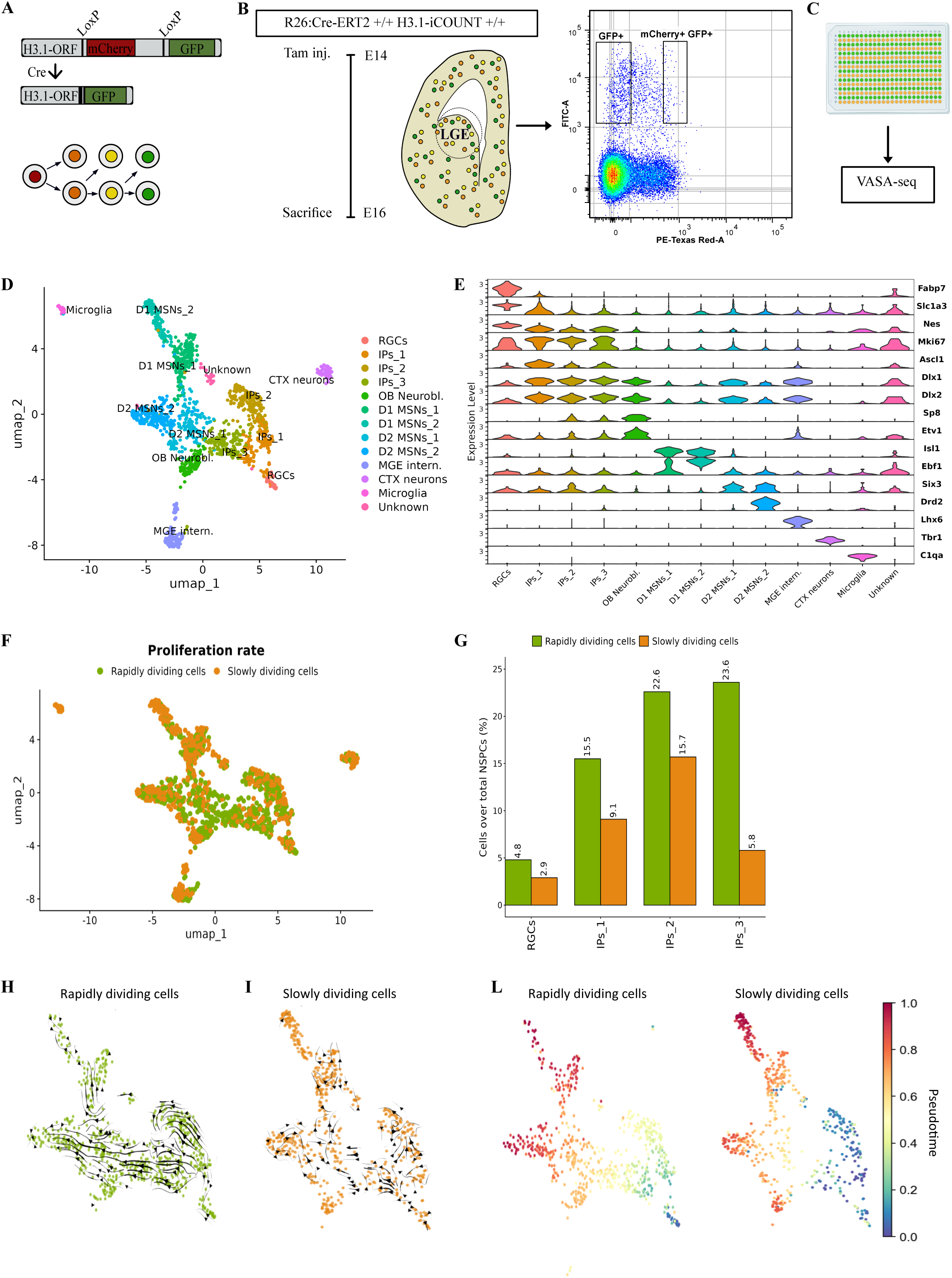
Isolation and transcriptomic profiling of slowly and rapidly dividing LGE cells. A. Illustration of the iCOUNT knock-in gene. B. Experimental paradigm used for tracing slowly and rapidly dividing NSPCs during the time window of aNSCs specification. Tamoxifen was injected into double homozygous pregnant females at E14 and embryonic LGEs were collected two days post injection (E16). Slowly (GFP+ mCherry+) and rapidly (GFP+ mCherry-) dividing cells were isolated via FACS. C. Illustration of sorting pattern used in B. Rapidly and slowly dividing cells were sorted in the same 384-well plates to limit batch effects during RNA-seq. Image of the 384-well plate was created with BioRender.com D. Uniform Manifold Approximation and Projection (UMAP) plot of the sorted cell populations. E. Violin plot showing normalized expression of known markers genes. F. UMAP plot depicting the proliferation rate of sorted cells. G. Histogram showing the proportion of slowly and rapidly dividing cells in each NSPC cluster over the total NSPCs. H-I. RNA velocity analysis of rapidly and slowly dividing cells. Arrow direction indicates the inferred trajectory of cellular differentiation or gene expression change; arrow length reflects the corresponding speed of change. L. Palantir-based pseudotime analysis of rapidly and slowly dividing cells. Cells are color-coded by pseudotime, with minimum and maximum values corresponding to the least differentiated and terminally differentiated states, respectively.

Count matrices from a total of 5 384-well plates – representing 3 biological replicates – were integrated using Seurat v5 (Fig. S1A). After stringent quality control (Fig. S1B-C), we retained 1,442 high-quality cells that were grouped into transcriptomic similar cell populations via unsupervised clustering analysis (Fig. 1D). To prevent clustering from being dominated by cell-cycle genes, we calculated a cell-cycle phase score using a list of canonical cell-cycle marker genes (Tirosh et al., 2016) and differences in G2M- and S-phase scores were regressed out during pre-processing (Fig. S1D). Importantly, this partial cell cycle regression preserves signals separating cycling and not cycling cells, thereby not affecting the identification of quiescent cells. Next, cells were classified into 13 clusters based on the expression of canonical markers (Fig. 1D-E). RGCs were classified as cells expressing *Fabp7*, *Slc1a3* and *Nestin;* IPs as cells expressing *Ascl1*, *Dlx1, Dlx2* and *Mki67*; OB neuroblasts were defined by the expression of *Dlx1, Etv1* and *Sp8;* finally, striatal medium spiny neurons (MSNs) were defined as cells expressing *Isl1* and *Ebf1* (direct pathway, D1 MSNs), and *Six3* and *Drd2* (indirect pathway, D2 MSNs). In addition, we detected a small contamination from the neocortex (*Tbr1*), medial ganglionic eminence (*Lhx6*), and microglia (*C1qa*), as well as one small cluster of unidentified cells likely representing low-quality cells that passed the quality control and characterized by lower counts and fewer genes detected per cell (Fig. S1E).

To identify slowly and rapidly dividing cells, we inferred the proliferation rate of the cells in our dataset by combining well-specific barcodes with defined sorting patterns (Fig. 1C, F). Notably, the *Tbr1*-expressing population was mainly derived from cells that divided a few times between E14 and E16 (Fig. S1G), consistent with the cell-cycle dynamics in the neocortex, where neurons are generated by NSPCs that divide less frequently than those in the ventral telencephalon. This data further supports the validity of the iCOUNT labelling strategy. Interestingly, among the NSPCs – comprising RGCs along with IPs_1, IPs_2, and IPs_3 – slowly dividing cells were almost equally abundant as the rapidly dividing counterpart, with the exception of cluster IPs_3 where they represented only ∼20% of the cell population (Fig. 1G). The presence of slowly dividing cells in different NSPC clusters might suggest different developmental origins of aNSCs. We therefore analysed more in detail the lineage progression of slowly and rapidly NSCPs by performing RNA velocity (La Manno et al., 2018). Consistent with their role as the main drivers of embryonic neurogenesis, rapidly dividing NSPCs displayed a clear differentiation trajectory originating from RGCs, progressing through IPs, and terminating in neuronal clusters (Fig. 1H). On the contrary, RNA velocity vectors were much reduced in length and more chaotic in the slowly dividing NSPCs, suggesting that differentiation proceeds at lower rate in this population (Fig. 1I). In agreement with this result, Palantir-based pseudotime analysis (Setty et al., 2019) showed that slowly dividing NSPCs exhibit lower pseudotime values, consistent with a less differentiated cellular state (Fig. 1L). Notably, slowly dividing RGCs displayed little dynamism and vectors pointing away from the IP clusters, suggesting that – contrary to rapidly dividing RGCs – these cells are not actively committing to an IP fate (Fig. 1H-I). On the other hand, similarly to their rapidly dividing counterparts, slowly dividing IPs displayed RNA velocity vectors directed toward more differentiated progenitors and neuronal clusters, suggesting a progression toward the neurogenic lineage. CellRank fate probability analysis (Lange et al., 2022) further supported this observation, revealing similar probabilities of reaching the neuronal terminal state in both slowly and rapidly dividing IPs (Fig. S1G). Interestingly, slowly dividing IPs_3 cells displayed also RNA velocity vectors with circular orientation, consistent with active cell cycling and possibly indicating amplification of the committed progenitor pool (Fig. 1I). Thus, while slowly and rapidly diving RGCs seem to diverge in their differentiation trajectory, IPs clusters are committed to the neurogenic lineage regardless of division rate.

### Distinct gene signatures in slowly dividing NSPCs converge on common biological processes

Given the presence of slowly dividing cells in both RGCs and IPs, we next asked whether the proliferative behaviour in these subpopulations was driven by similar molecular mechanisms. To address this point, we performed differential gene expression (DE) analysis between rapidly and slowly dividing cells in each stem/progenitor cluster and compared the transcriptomic signatures among NSPCs (Fig. 2A-D). Interestingly, this analysis revealed no common genes enriched among slowly dividing cells in different NSPC clusters (Fig. 2E). However, using Gene Ontology analysis we identified a set of biological processes shared among different slowly dividing NSPC populations, suggesting that distinct genes converge on similar molecular pathways (Fig. 2F-I). For instance, slowly dividing RGCs and IPs_1 showed enrichment of genes associated with protein-DNA complex organization, nucleosome assembly and epigenetic regulation (e.g. *Atad2*, *Chd2*, *Suz12*, *Dnmt1*, *Terf1*, *Med25*), which suggest extensive chromatin remodelling activity in these two populations (Fig. 2F-G). Of note, slowly dividing RGCs also displayed genes associated with cell adhesion (e.g. *Cdh11*, *Cd9*, *Cd81*, *Lamb2, Fbln2*, *Fn1*), similarly to what has been previously reported in quiescent NSCs (Llorens-Bobadilla et al., 2015; Morizur et al., 2018). In contrast, genes enriched in slowly dividing IPs_2 and IPs_3 were associated with ribosome biogenesis (e.g. *Rpl7a*, *Rpl7l1*, *Wdr12*, *Wdr18*, *Tbl3*) and c-Jun N-terminal kinase cascade (e.g. *Mink1*, *Dvl2*, *Traf4*, *Daxx*, *Tlr4*, *Mapk8ip2*) (Fig. 2H-I), processes linked to NSPCs proliferation and differentiation (Baser et al., 2017; Kim et al., 2007). The enrichment of genes associated with both proliferation and differentiation is consistent with the RNA velocity analysis, suggesting that slowly dividing IPs_2 and IPs_3 are primed for differentiation while retaining proliferative capacity, thus likely representing cells capable of generating additional committed progenitors.

**Figure 2.**
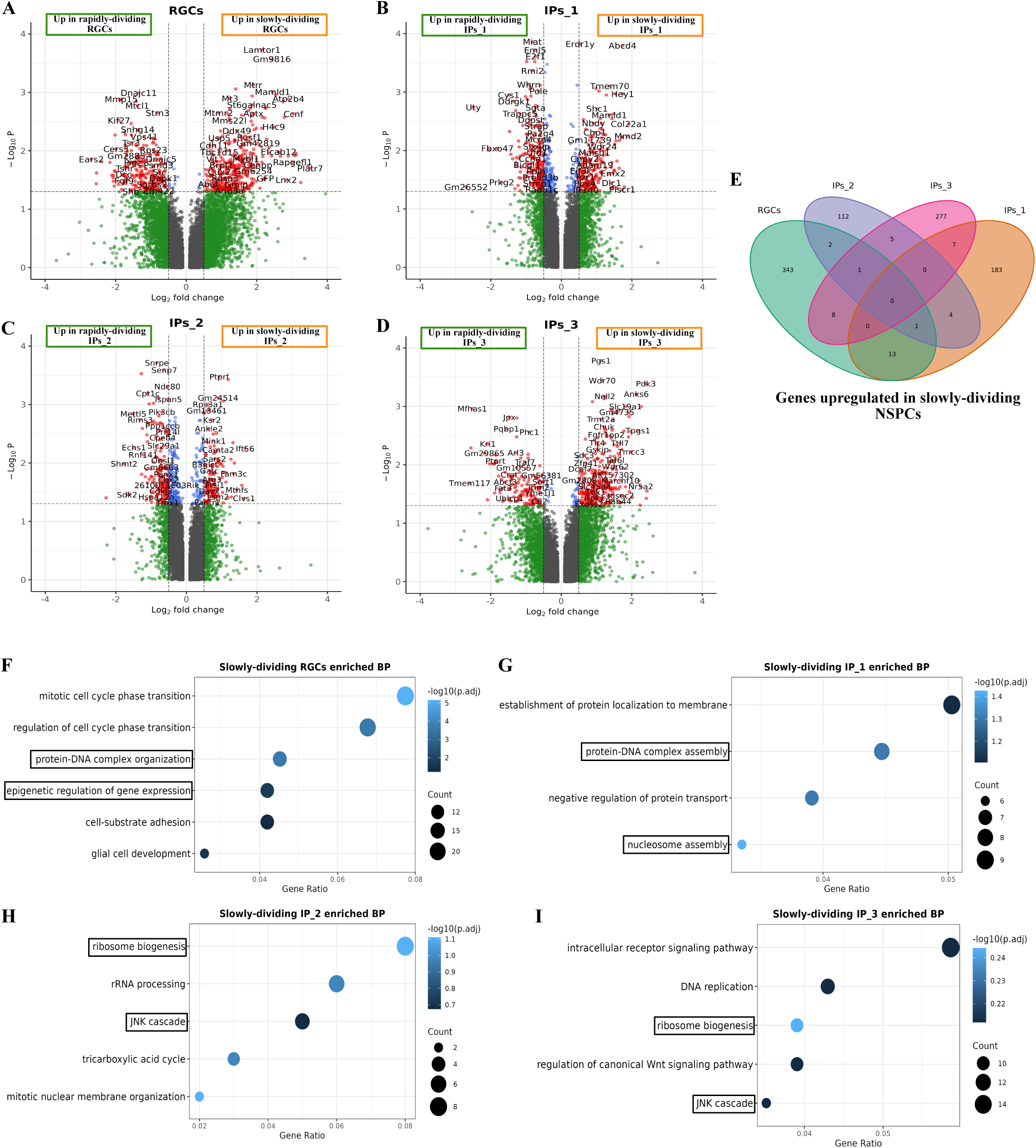
Differential gene expression analysis between slowly and rapidly dividing LGE NSPCs. A-D. Volcano plots showing differentially expressed genes (DEGs) between slowly and rapidly cells in RGC and IP clusters. E. Venn diagram showing overlap in slowly dividing cells-enriched DEGs across RGC and IP clusters. F-I. Selected biological processes (BP) enriched in slowly dividing RGC and IP clusters.

Similarly to slowly dividing NSPCs, rapidly dividing stem/progenitor cells showed distinct transcriptomes across clusters but falling into similar biological processes, predominantly associated with cell proliferation (Fig. S2). Interestingly, some of these proliferation-associated processes (i.e. DNA replication, ribosome biogenesis and WNT signalling pathway) were also enriched in the slowly dividing IPs_2 and IPs_3 clusters (Fig. 2H-I and Fig. S2A-C), suggesting that these cells may be transitioning toward a more proliferative state.

### Overlap in gene expression signatures between slowly dividing RGCs and postnatal neural stem cells

Adult NSCs share several hallmarks with embryonic RGCs, such as apico-basal polarity, self-renewal capacity and expression of astroglia markers (e.g. *Blbp*, *Slc1a3*, *Aldoc*, *Aldh1l1)*. In addition, our transcriptomic analysis highlighted the enrichment of cell adhesion-related genes in slowly versus rapidly dividing RGCs (Fig. 2F), a transcriptional feature previously described in quiescent NSCs (Llorens-Bobadilla et al., 2015; Morizur et al., 2018). We therefore explored the population of slowly dividing RGCs more in detail. To identify the mechanisms underlying the emergence of aNSCs, we compared the genes enriched in slowly dividing RGCs with the ones expressed in postnatal NSCs (pNSCs), the latter obtained from a published dataset of the lateral V-SVZ at postnatal day (P) 29-35 (Cebrian-Silla et al., 2021) (Fig. 3A). The gene signature of pNSCs was created by performing DE analysis of pNSCs versus transit-amplifying progenitors and neuroblasts, which form the rest of the neurogenic lineage in the V-SVZ (Fig. S3A). By overlapping the gene lists from pNSCs and slowly dividing RGCs we identified a set of 62 shared genes (Fig. 3B). Many of these were associated with different metabolic processes, including biosynthesis of fatty acids which have been shown to regulate NSCs proliferation in the V-SVZ (Stoll et al., 2015). Interestingly, slowly dividing RGCs and pNSCs also showed enrichment of genes involved in cell-substrate adhesion (Fig. 2F, 3C), suggesting that niche interactions might be a common mechanism regulating cell behaviour in both populations. Of note, the extracellular environment is a key regulator of NSC proliferation and quiescence in the adult V-SVZ (Kazanis et al., 2010; Lim et al., 2000; Porlan et al., 2014), where quiescent NSCs are known to express several ECM and cell adhesion molecules (Hu et al., 2017; Kokovay et al., 2012; Llorens-Bobadilla et al., 2015; Morizur et al., 2018). Similarly to the adult neurogenic niche, the extracellular environment might regulate the balance between rapidly dividing neurogenic and slowly dividing NSPCs in the embryonic neurogenic niche. A detailed analysis of the common gene signature among pNSCs and slowly dividing RGCs revealed the ∼20% of the shared genes were associated to ECM remodelling (*Lamb2*, *Timp3, Fbln2, Vit, Axl, Gfod2)* or cell adhesion and signalling (*Cdh11*, *Trip6, Cd9*, *Cd81, Gpc6, Sirpa, Egr1*) (Fig. S3B-C). Interestingly, among the most upregulated genes in rapidly dividing RGCs, we identified *Mmp15* (Fig. 2A, 3D), a member of the family of matrix metalloproteinases (MMPs), enzymes involved in the degradation of ECM molecules such as laminins and collagens (Itoh, 2015).

**Figure 3.**
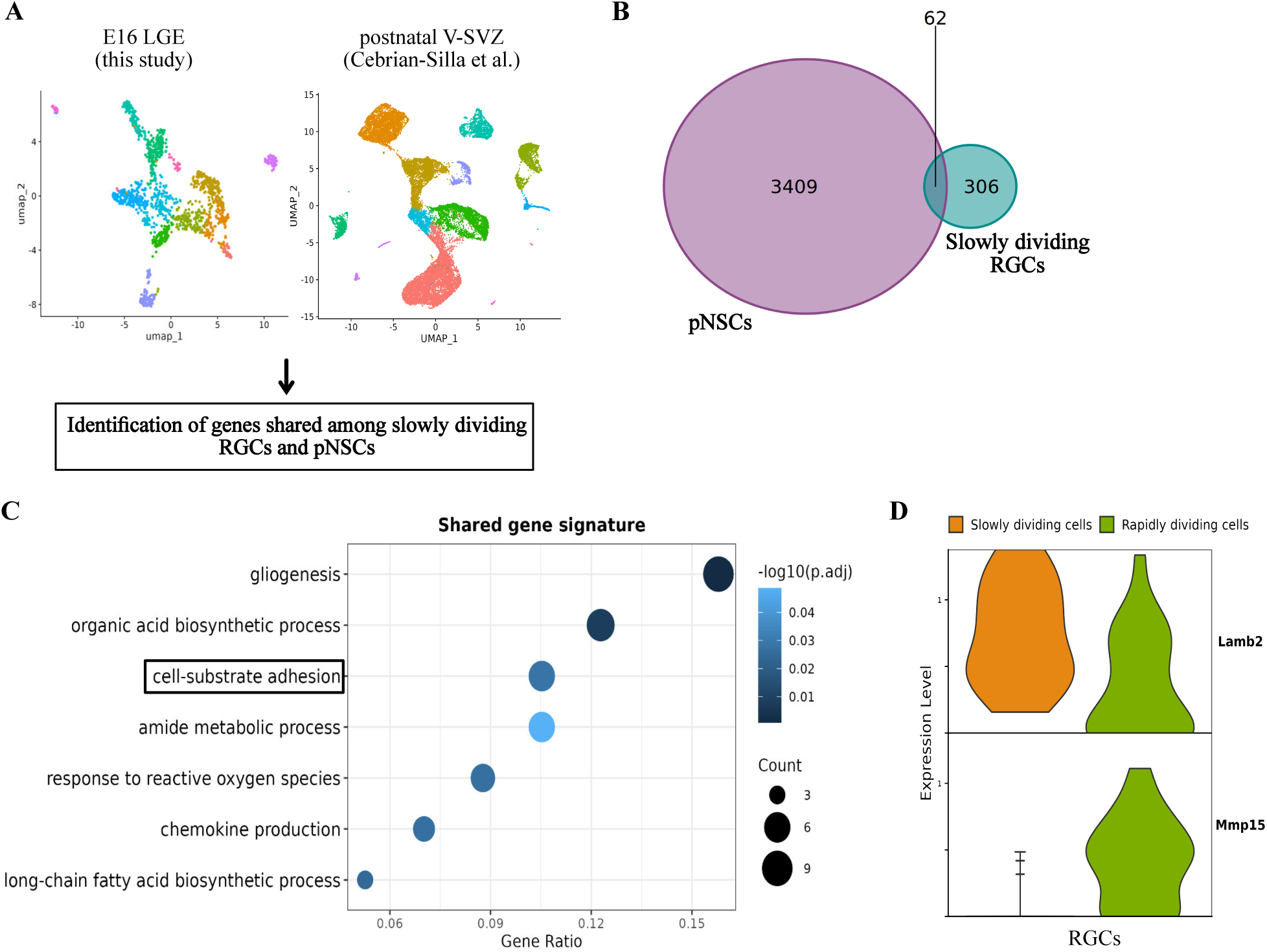
Identification of gene signature shared among slowly dividing RGCs and postnatal NSCs and selected candidate genes. A. Workflow used to identify common gene signature between slowly dividing RGCs (this study) and postnatal NSCs (Cebrian-Silla et *al.*, 2021). B. Venn diagram showing overlap in the genes enriched in postnatal NSCs (pNSCs) versus TAPs and neuroblasts and in genes enriched in slowly versus rapidly dividing RGCs. C. Selected biological processes enriched in pNSCs and slowly dividing RGCs. D. Violin plot showing normalized expression of selected candidate genes.

These observations prompted us to investigate the contribution of the extracellular environment to the specification of aNSCs. To do so, we explored in more detail the roles of two ECM-related genes, *Lamb2* and *Mmp15*, enriched in slowly and rapidly dividing RGCs, respectively (Fig. 3D). *Lamb2* encodes the ꞵ2-chain of laminins, heterotrimeric proteins secreted in the extracellular space and formed by the interaction of α, ꞵ and γ chains, which are present in different isoforms. LAMB2 has been previously characterized as a component of the adult neurogenic niche where it is part of specific ECM-structures *–* known as *fractones –* (Sato et al., 2019) and known to modulate proliferation in this region (Nascimento et al., 2018). *Mmp15* encodes a MMP previously implicated in cell migration (Tao et al., 2011). Differently from other MMPs secreted in the extracellular space, MMP15 is part of the sub-family of membrane-bound MMPs, characterized by a transmembrane domain, which might limit their action to the pericellular area (Itoh, 2015). Thus, while slowly dividing RGCs express genes encoding ECM components – such as *Lamb2* – rapidly dividing RGCs might promote ECM turnover in their vicinity through expression of *Mmp15*.

### Expression profile of *Lamb2* and *Mmp15* in the embryonic telencephalon

To investigate the role of *Lamb2* and *Mmp15* in the emergence of aNSCs, we first analysed their expression profiles in the embryonic neurogenic niches at different developmental time points. In situ hybridization analysis revealed a dynamic expression pattern of *Lamb2*. Indeed, even though enriched in the LGE germinal zones at all the stages analysed, *Lamb2* expression became gradually enriched at the most apical surface of the VZ by E16, i.e. the end of the time-window of aNSCs specification (Fig. 4A). On the contrary, *Mmp15* was broadly expressed across the germinal zones throughout the telencephalon, at all the time points analysed (Fig. 4A). This is in line with our scRNA-seq data showing expression of *Mmp15* in all the NSPC clusters, as well as in the neuronal populations, while *Lamb2* expression is restricted mainly to the RGC and IPs_1 clusters (Fig. 4B-C). Notably, *Lamb2* and *Lama5* were the only members of the laminin family showing such an enrichment in RGCs and IPs_1, as compared to other clusters (Fig. S4).

**Figure 4.**
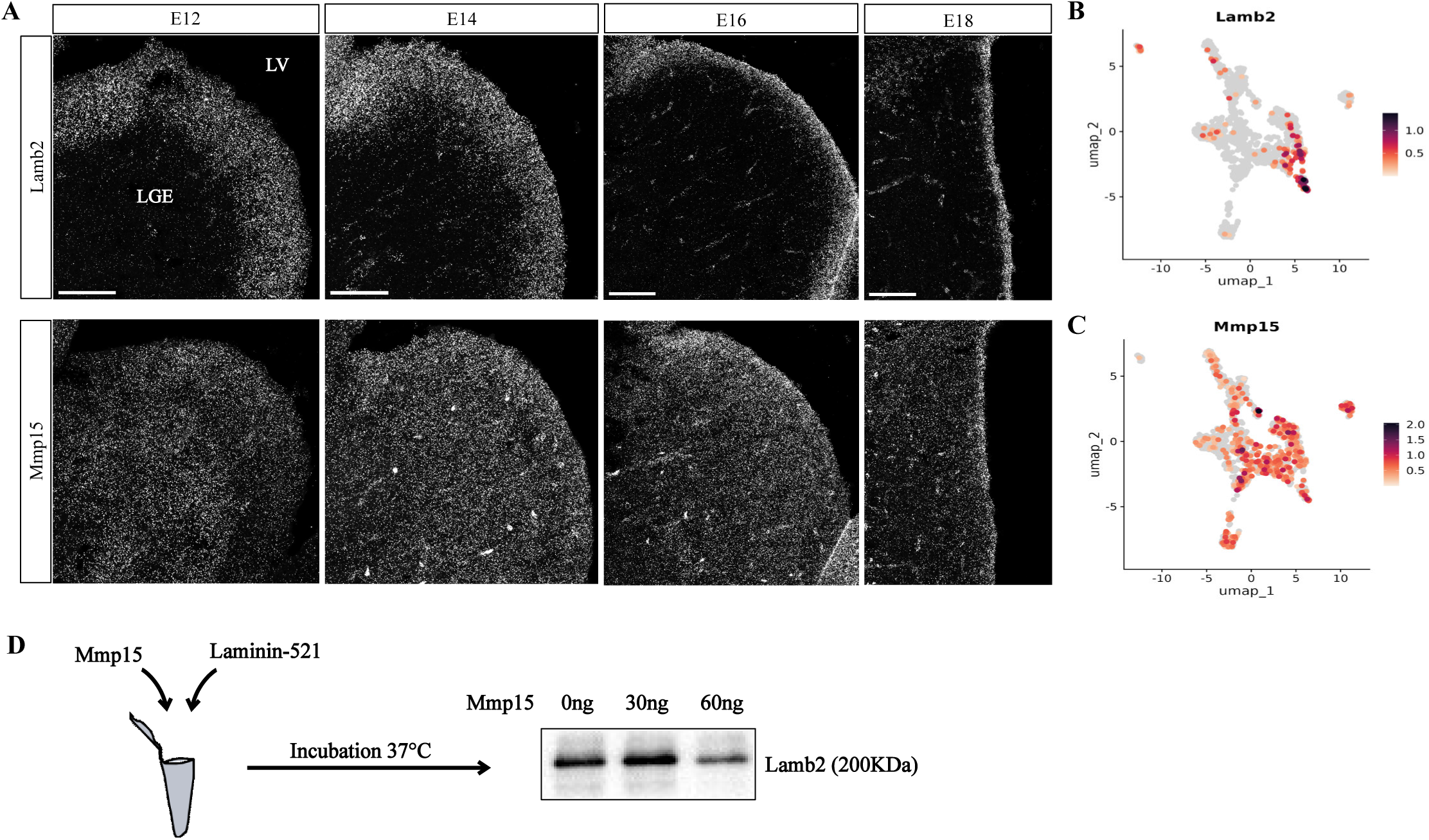
Expression pattern of *Lamb2* and *Mmp15* during development and *in vitro* degradation assay for MMP15 and LAMB2. A. RNA scope showing expression of *Lamb2* and *Mmp15* in embryonic LGE at the different developmental stages indicated. Note that *Lamb2* expression is restricted to the apical VZ by E16, while *Mmp15* is expressed across the entire LGE at all the time points analyzed. B-C. UMAP showing normalized expression of *Lamb2* and *Mmp15* across clusters. Note that *Lamb2* is mainly expressed by RGCs and IPs_1, while *Mmp15* is expressed in both NSPC and neuronal clusters. D. *In vitro* degradation assay for MMP15 and LAMB2. Different concentrations of purified activated MMP15 were incubated 4 hours at 37°C with purified β2-containing laminin (Laminin-521). Western blot analysis showed reduced LAMB2 band intensity in presence of 60ng of MMP15.

These results suggest a distinct remodelling of the ECM by slowly and rapidly dividing RGCs, where the former seems to initiate building the “adult” neurogenic niche via localized secretion of specific molecules, such as LAMB2, which in turn might promote the maintenance of the stem-cell fate, while rapidly-dividing RGCs and other NSPCs might “escape” such a microenvironment by upregulating proteases, such as MMP15, and progress toward the neurogenic lineage. To further test this hypothesis, we performed an *in vitro* degradation assay to assess the ability of MMP15 to degrade LAMB2. Different concentrations of purified MMP15 recombinant protein were incubated with a purified recombinant laminin-521 – containing the β-2 chain, in combination with the α-5 and γ-1 chains – and the effects on LAMB2 protein were measured via western blot (Fig. 4D). We observed a decrease in the intensity of the LAMB2 band which was inversely correlated with the amount of MMP15 protein present (Fig. 4D), thus validating the degradation capacity of MMP15 versus LAMB2.

### *Lamb2* and *Mmp15* play opposite roles in the specification/maintenance of postnatal NSCs

We then asked whether *Lamb2* and *Mmp15* could play a role in the specification and maintenance of postnatal NSCs (pNSCs). To address this point, we designed microRNAs versus our target genes to downregulate their expression (Fig. S5A) and cloned them together with a *GFP* reporter into lentiviral vectors for *in vivo* loss of function analysis. Knockdown (KD) of *Lamb2* and *Mmp15* was carried out at E13 and infected GFP+ cells in the lateral V-SVZ were analysed at P10 (Fig. 5A). Specifically, as pNSCs originate from slowly dividing cells, we injected the synthetic thymidine analogue 5-bromo-2′-deoxyuridine (BrdU) into the pregnant female at E13 to label cycling cells (Fig. 5A). We then analysed the slowly dividing BrdU-labelling retaining cells in the postnatal lateral V-SVZ showing NSCs features, such as a radial morphology and immunostaining for the astroglia marker GFAP. Interestingly, among all the infected cells, the percentage of radial GFP+ GFAP+ BrdU-retaining cells was reduced following *Lamb2* KD (Fig. 5B-C). *Lamb2* has been shown to regulate cell-cycle dynamics in retinal progenitor cells, via modulation of cell-cycle exit and mitotic spindle orientation (Serjanov et al., 2018; Serjanov et al., 2022). To test whether the reduction in the pNSCs pool following *Lamb2* KD was due to altered cell divisions, we performed immunohistochemical analysis for the proliferation marker KI67. This analysis revealed an overall increase in dividing cells in the postnatal V-SVZ (Fig. S5B-C). This is consistent with the study of Nascimento and colleagues showing that deletion of a distinct laminin chain isoform, *Lama5*, reduces the number of slowly cycling cells and increases proliferation in the postnatal V-SVZ (Nascimento et al., 2018), further reinforcing the role of laminins in modulating cell division in the V-SVZ. We then asked whether downregulation of *Mmp15* in rapidly dividing RGCs would instead promote the emergence of cells with NSC features. To this end, we applied the same labelling paradigm mentioned above and induced *Mmp15* KD at E13. Notably, *Mmp15* KD induced the opposite phenotype, namely an increase of radial GFP+ GFAP+ BrdU-retaining cells as compared to control (Fig. 5D-E). Importantly, the increase in radial GFP+ GFAP+ cells retaining BrdU label indicates that downregulation of the metallo-protease reduces the proliferation rate and promotes a quiescent NSC-like state in NSPCs at E13, when BrdU is injected.

**Figure 5.**
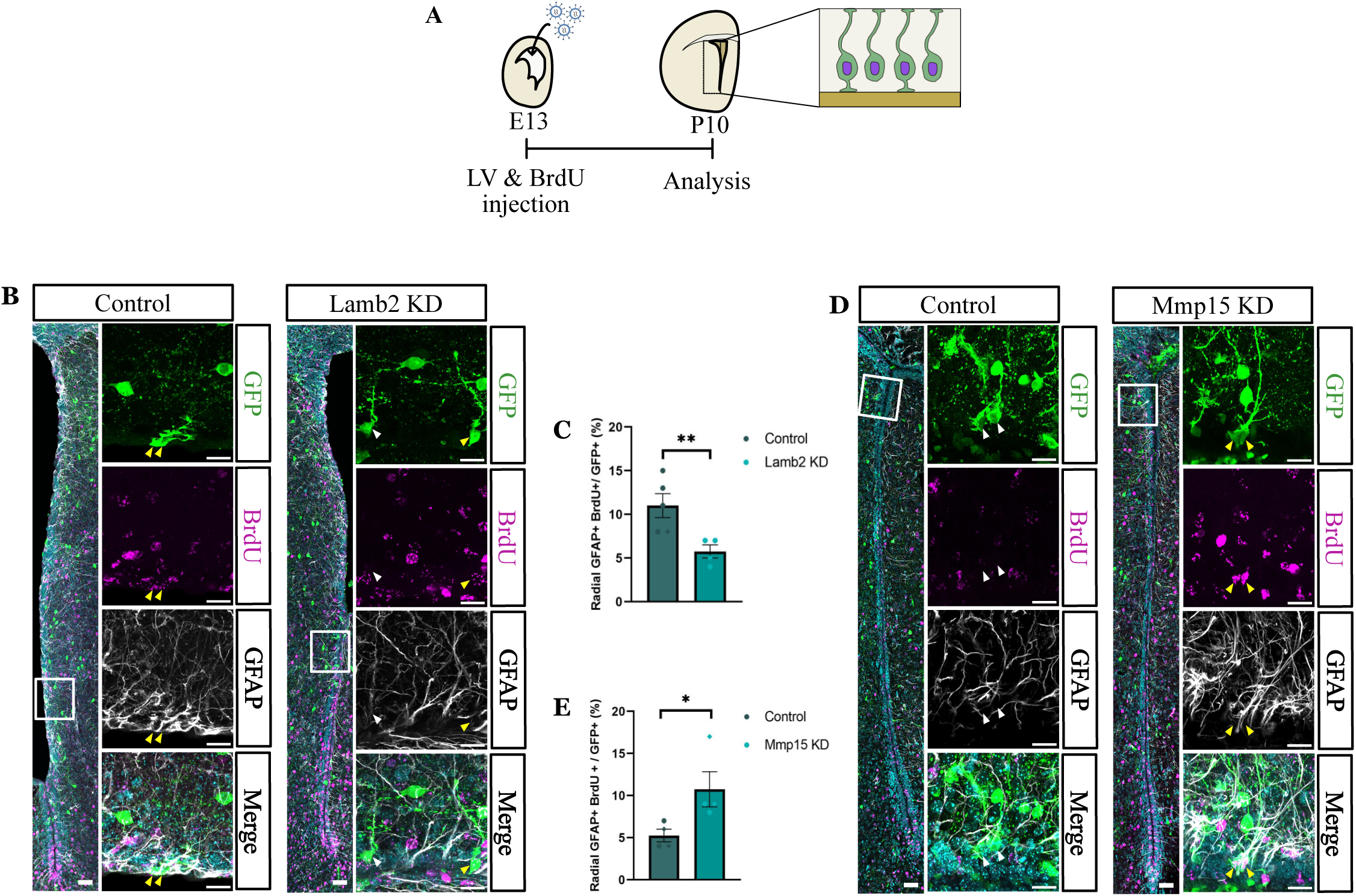
*In vivo Lamb2* and *Mmp15* KDs affect the number of radial GFP+ GFAP+ BrdU-retaining cells in the postnatal lateral V-SVZ in opposite way. A. Schematic illustration of experimental paradigm for *Lamb2* and *Mmp15* KDs and BrdU label-retaining analysis. Lentiviral vectors carrying miRNAs against candidate genes or a non-targeting miRNA were injected in the lateral ventricles of embryos at E13. After in utero injection (IUI), BrdU was injected into the pregnant females to label dividing cells in the litter. Radial GFP+ GFAP+ BrdU-labeling retaining cells were quantified in the lateral V-SVZ of P10 pups. LV: lentivirus. B, D. Representative images of the quantification shown in C, E. Yellow and white arrows indicate radial GFP+ GFAP+ cells with and without BrdU-label retention, respectively. Scale bar: 50μm in the overview images, 20μm in high-magnification images. C, E. Quantification of the percentage of radial GFP+ GFAP+ BrdU-retaining cells among all GFP+ cells in the lateral V-SVZ. Data are shown as means ± SEM. Dots with different shapes correspond to pups from different litters. * p< 0.05 and ** p< 0.01 by Mann-Whitney Test.

### Molecular analysis reveals increase of NSCs and changes in progeny upon *Mmp15* knockdown

Even though label retention assay, together with GFAP expression and morphological radial glia-like hallmarks, have been widely used to identify NSCs *in vivo*, it is conceivable that only some hallmarks of NSCs are elicited upon manipulating MMP15. Therefore, we set out to examine the NSC identity of the infected cells in the postnatal V-SVZ by performing single-cell RNA sequencing (scRNA-seq) in the condition where we observed increased percentage of cells displaying NSC-hallmarks, namely following *Mmp15* KD. Using the same experimental paradigm as before, we injected lentiviral vectors carrying a microRNA versus *Mmp15* or a non-targeting microRNA into the lateral ventricles of E13 embryos and collected the lateral V-SVZ at P10 (Fig. 6A). Importantly, to assess whether the knockdown affects also the surrounding cells in a cell non-autonomous manner, we profiled the transcriptome of both GFP+ and GFP− cells (Fig. S6A-B). A total of 12341 cells that passed the quality control were classified into ependymal cells (*Foxj1*, *S100b*), NSCs (*Gfap*, *Sox9*, *Fabp7*, *Slc1a3*, *Aldh1l1*, *Aldoc*, *Hes5*, *Vcam1*), neurogenic (*Ascl1*, *Dcx*, *Dlx1*, *Sp8*, *Sp9*) and gliogenic (*Ascl1*, *Sox10*, *Pdgfra*, *Olig2*) transit-amplifying progenitors (TAPs), neuroblasts (*Dcx*, *Dlx1*, *Sp9*, *Sp8*, *Etv1*), oligodendrocytes (*Sox10*, *Olig2*, *Mbp*) and microglia (*C1qa*) (Fig. 6B-C). Notably, all the 3 clusters of NSCs express the quiescence marker *Vcam1* (Hu et al., 2017; Kokovay et al., 2012), but the different expression levels of *Egfr,* a gene associated with NSCs activation (Codega et al., 2014; Llorens-Bobadilla et al., 2015), suggest that the identified NSC populations might be in distinct quiescence states, with cluster NSCs_2 primed for activation (Fig. 6C). Interestingly, in line with the immunohistochemical analysis, we observed more than 2- and 3-fold increase in NSCs_1 and NSCs_3, respectively, following *Mmp15* KD, and a milder increase in NSCs_2 (Fig. 6C). These data suggest that *Mmp15* KD does not lead to higher expression levels of GFAP, or aberrant cell populations, but rather increases the NSC lineage. This conclusion is further corroborated by the increase in the proportion of ependymal cells, which share a common developmental origin with aNSCs (Ortiz-Álvarez et al., 2019; Redmond et al., 2019), suggesting that *Mmp15* KD enlarges the pool of a common aNSC and ependymal cell ancestor. Similar results were obtained when comparing cell proportion among not infected cells (Fig. S6C), highlighting *Mmp15*-mediated cell non-autonomous mechanisms on the neighboring cells. NSCs have been shown to give rise to both glial and neuronal cells (Figueres-Oñate et al., 2019; Mizrak et al., 2019). However, oligodendrocytes were not affected in numbers by *Mmp15* KD, while *Dcx*-expressing neuroblasts were decreased as compared to control. This could be due to NSCs entering quiescence and consequently ceasing neuroblast production, with the absence of an effect on oligodendroglial lineage pointing toward the existence of separate pools of neurogenic and gliogenic NSCs, as previously shown in the dorsal V-SVZ (Ortega et al., 2013).

**Figure 6.**
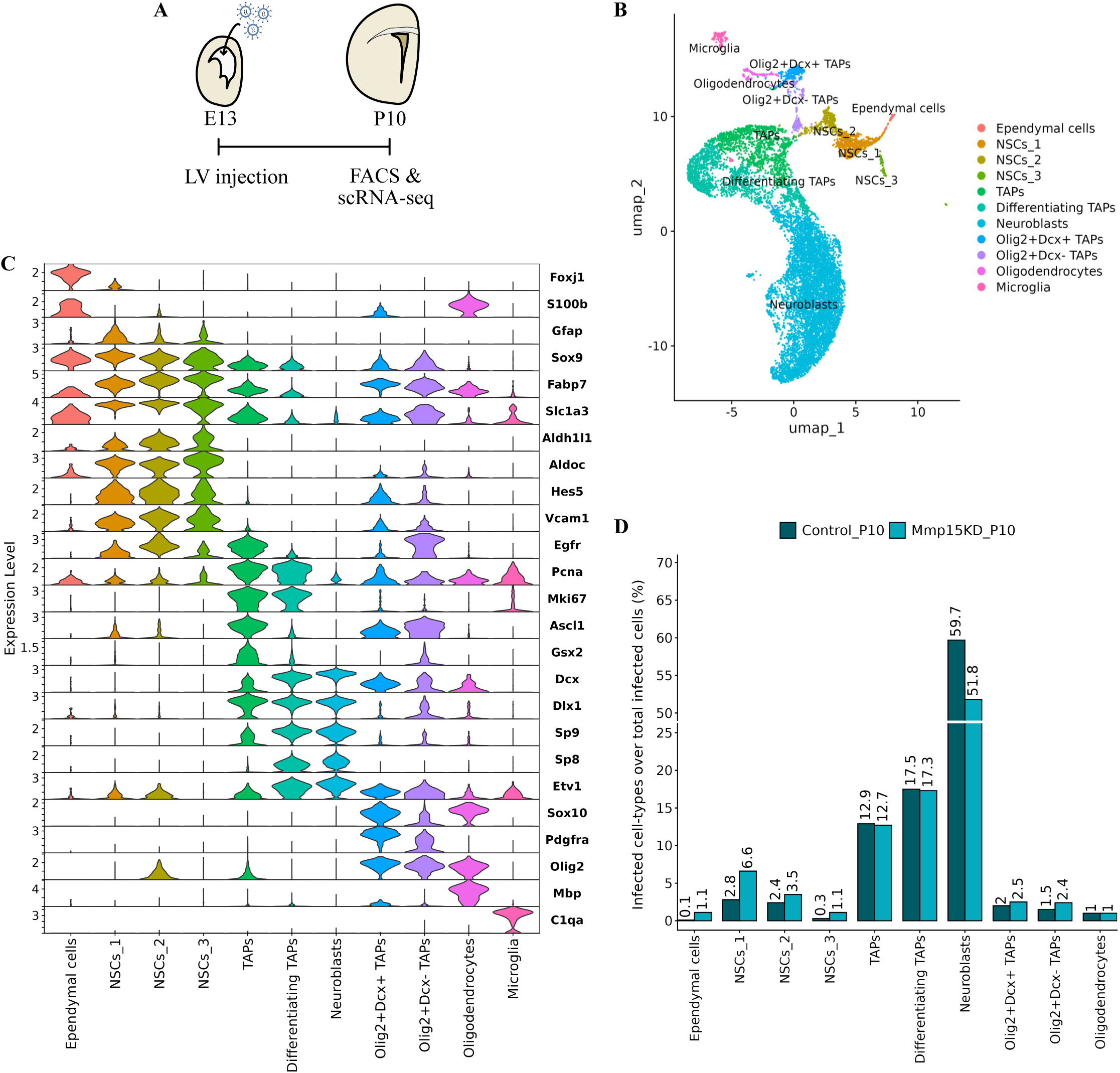
Transcriptome of P10 V-SVZ cells following *Mmp15* KD. A. Experimental paradigm used. B. UMAP plot of merged *Mmp15* KD and Control datasets showing identified cell populations composed of both infected and not infected cells. C. Violin Plot showing normalized expression of known marker genes labeling the main cell populations in the postnatal V-SVZ. D. Histogram showing the proportion of infected cell-types over all the infected cells. Infected cells were identified using the viral Woodchuck Hepatitis Virus Posttranscriptional Regulatory Element (WPRE) sequences (see Materials and Methods for details).

### *Lamb2* and *Mmp15* differentially affect gliogenesis and neurogenesis in the postnatal V-SVZ

In light of these results, we sought to examine the effects of *Mmp15* and *Lamb2* KDs on both the gliogenic and neurogenic lineages in the lateral V-SVZ via immunohistochemical analysis using the glial marker *Olig2* or the neuroblast marker *Dcx*. Consistent with scRNA-seq results, *Mmp15* KD showed significant effects only on the neurogenic lineage with a lower percentage of GFP+ DCX+ cells as compared to control (Fig. 7A-D). The reduced generation of neuroblasts following *Mmp15* KD might be a consequence of increased quiescence of pNSCs, as shown by increased percentage of BrdU-labelling retaining pNSCs (Fig. 5D-E).

**Figure 7.**
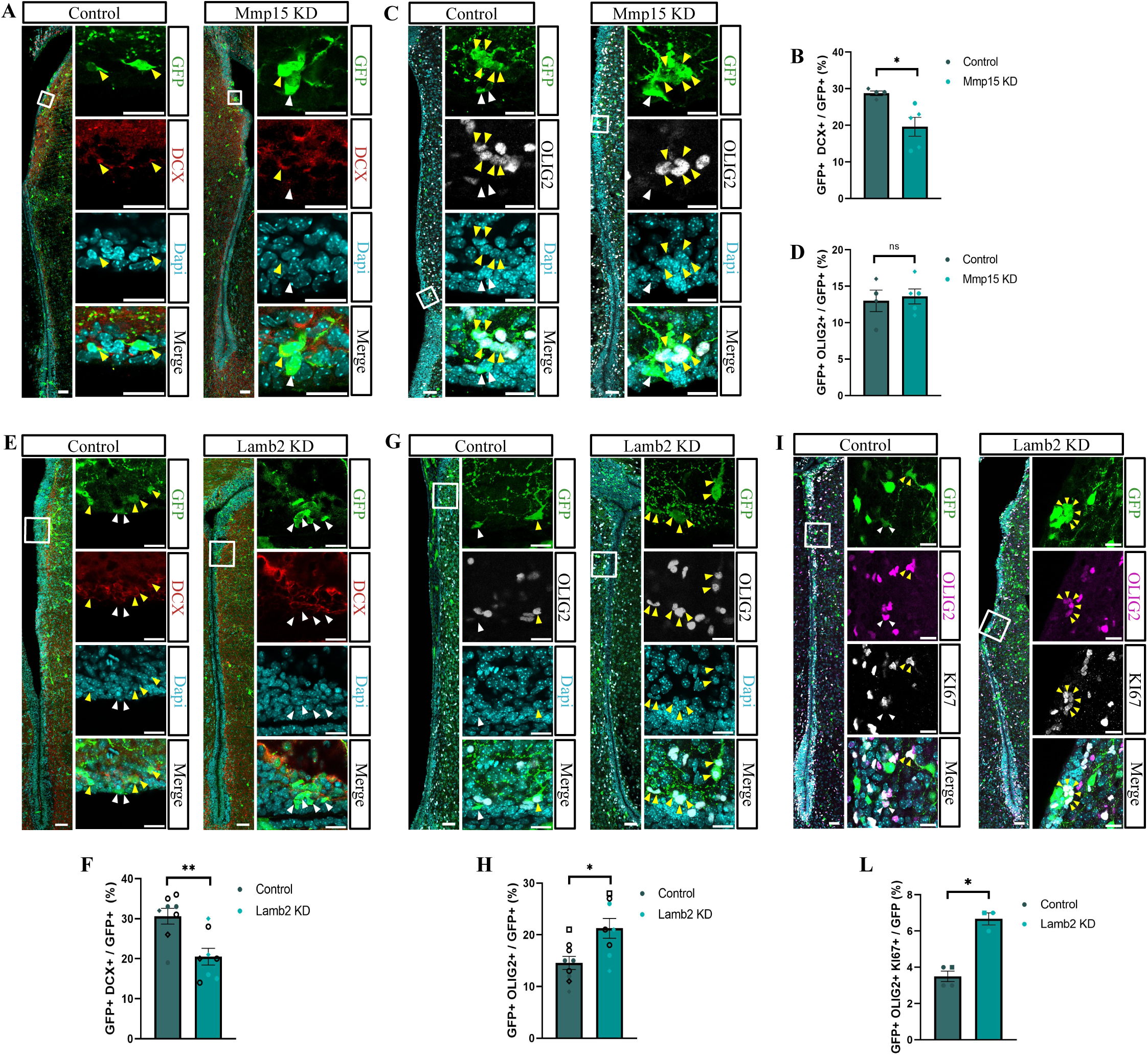
*In vivo Lamb2* and *Mmp15* KDs reduce neurogenesis and differentially impact gliogenesis in the postnatal lateral V-SVZ. A-D. Representative images and quantification showing reduction of GFP+ DCX+ cells (A-B) and no effects of GFP+ OLIG2+ cells (C-D) in the P10 lateral V-SVZ following *Mmp15* KD. Data are shown as means ± SEM. Dots with different shapes correspond to pups from different litters. E-I. Representative images and quantification showing reduction of GFP+ DCX+ cells (E-F) and increase in GFP+ OLIG2+ cells (G-H) and GFP+ OLIG2+ KI67+ cells (I-L) in the P10 lateral V-SVZ following *Lamb2* KD. Data are shown as means ± SEM. Filled and empty dots represent data from lentivirus- and retrovirus-mediated KD, respectively. ns: not significant, * p< 0.05 and ** p< 0.01 by Mann-Whitney Test. Yellow and white arrows indicate cells positive or negative for shown markers, respectively. Scale bar: 50μm in the overview images, 20μm in high-magnification images.

Conversely, *Lamb2* KD affected both glial and neuronal production, showing a significant decrease in GFP+ DCX+ neuroblasts (Fig. 7E-F), while GFP+ OLIG2+ cells were increased amongst the KD cells (Fig. 7G-H). Considering the role of *Lamb2* in cell proliferation (Fig. S5B-C), we tested whether the increase in glial cells was due to increased proliferation. Immunohistochemical analysis showed indeed a higher percentage of dividing GFP+ KI67+ OLIG2+ cells in the KD condition as compared to control (Fig. 7I-L). Interestingly, we also observed many KI67-only-positive cells with small nuclei reminiscent of TAPs (data not shown), suggesting that cells might get trapped in an amplifying state and fail to progress to neuroblasts. Altogether, these data suggest that manipulation of the extracellular environment via distinct ECM-associated proteins differentially affects the emergence of pNSCs as well as gliogenesis and neurogenesis in the V-SVZ.

## Discussion

Here, we show that slowly dividing cells in the embryonic LGE are a heterogeneous population comprising both RGCs and IPs and characterized by different gene signatures. The presence of slowly dividing cells across distinct NSPC clusters may point to different developmental origins of aNSCs. Even though we cannot exclude this, our data suggest that the slowly dividing cells present in the IP clusters might have already initiated the differentiation program, as shown by the expression of *Dlx1* and *Dlx2,* which are required for neuronal differentiation (Anderson et al., 1997), as well as by the RNA velocity analysis showing IP cells progressing toward the neuronal clusters, a finding supported also by CellRank fate probability analysis. Moreover, genes enriched in slowly dividing IPs_2 and IPs_3 are associated with processes such as ribosome biogenesis and JNK signalling, which have been linked to NSCs activation, proliferation and differentiation (Castro-Torres et al., 2024; Kim et al., 2007; Llorens-Bobadilla et al., 2015). Therefore, these data suggest that the population of slowly dividing cells in IPs_2 and IPs_3 might be primed for differentiation but still be able to undergo cell division, likely being actively dividing cells that amplify the pool of committed progenitors. Along this line, previous live imaging experiments have shown that the cell cycle length of progenitors generated after one cell division is longer than the one of progenitors produced after multiple rounds of cell divisions, highlighting a shortening of the cell cycle duration at each progenitor generation (Pilz et al., 2013). Consequently, slowly dividing cells present in the committed IP clusters might represent the first generation of progenitors produced between E14-E16, retaining proliferative capacity – and likely being only temporary “slowly dividing”–and giving rise to other progenitors, rather than being ancestors of aNSCs.

Focusing on RGCs, we found that slowly dividing RGCs and pNSCs share gene sets associated with ECM-remodelling and cell adhesion. This is particular interesting as, in the adult neurogenic niche, molecules involved in cell-cell and cell-ECM interactions have been shown to be preferentially enriched in dormant NSCs, as compared to other niche cell populations, and modulate NSCs quiescence (Kokovay et al., 2012; Llorens-Bobadilla et al., 2015; Morizur et al., 2018; Porlan et al., 2014). Thus, the expression of several common niche-related genes suggests that slowly dividing RGCs have already acquired a key feature of quiescent aNSC and that similar niche-associated mechanisms might modulate the maintenance of these cell populations. This is of particular interest, as in the adult V-SVZ aNSCs are the only cell type whose expression mirrors the ECM proteome of the niche, suggesting that they create their own local niche by secretion of ECM molecules (Kjell et al., 2020). The expression of the same ECM-associated genes in both pNSCs and slowly dividing RGCs indicates that, already at early developmental stages, slowly dividing RGCs create a local microenvironment which is, at least in part, similar per composition to the adult neurogenic niche. We hypothesize that such a niche might in turn favour the maintenance of the stem cell identity. In line with this hypothesis, among the share genes, we identified *Timp3,* an inhibitor of the MMPs, previously shown to be enriched in slowly versus rapidly dividing NSPCs and suggested to promote the maintenance of an undifferentiated state in LGE NSPCs (Fang et al., 2023). Of note, TIMP3 can inhibit the protease activity of MMP15 (Butler et al., 1997) *–* which we found among the most upregulated genes in rapidly versus slowly dividing RGCs*–* and therefore it could limit the MMP15-mediated ECM turnover in the vicinity of slowly dividing RGCs, promoting the maintenance of the local microenvironment. This is particularly relevant considering that slowly dividing RGCs express LAMB2, which we showed being degraded by MMP15 *in vitro.* In this regard, it would be interesting to assess whether MMP15 is able to degrade other ECM proteins produced by slowly dividing RGCs. Overall, these data highlight the presence of distinct ECM-remodelling programs active in slowly and rapidly dividing RGCs.

To assess whether distinct ECM-remodelling mechanisms could regulate the developmental emergence of aNSCs, we performed *in vivo* loss-of-function analysis of *Mmp15* and *Lamb2* at E13, when aNSCs start to be set aside. In order to identify NSCs we used well-established aNSC-features. For instance, live imaging and immunohistochemical analysis have shown that NSCs in the postnatal and adult V-SVZ are characterized by a radial morphology, with a basal process contacting blood vessels and an apical process facing the lateral ventricle (Merkle et al., 2004; Obernier et al., 2018; Takemura et al., 2025) – the latter being absent in a recently identified subpopulation of basally located NSCs (Baur et al., 2022; Cebrian-Silla et al., 2025). Fate mapping experiments have demonstrated that most of the NSCs express the astroglia marker GFAP (Beckervordersandforth et al., 2010; Garcia et al., 2004), which is absent or lowly expressed in other niche cell populations such as transit-amplifying progenitors (TAPs) and ependymal cells (Doetsch et al., 1999). Finally, NSCs are derived from embryonic NSPCs that slow down their cell divisions between E13 and E16, thereby becoming identifiable through label-retention assays (Fuentealba et al., 2015; Furutachi et al., 2015). By using these key NSC features, we could assess the effects of the KDs on the NSC population, where we observed an increased percentage of NSC-like cells in the postnatal V-SVZ following *Mmp15* KD, and the opposite phenotype following *Lamb2* KD. Therefore, in physiological conditions *Lamb2* expression promotes the establishment of cells with aNSC hallmarks, while *Mmp15* plays the opposite role. In line with this, transcriptomic profiling of the *Mmp15* KD condition revealed an increased proportion of cells with an NSC identity, i.e. characterized by the expression of genes typically expressed by aNSCs.

LAMB2 has been previously described as a component of the adult neurogenic niche (Kjell et al., 2020; Sato et al., 2019). Specifically, Kjell et al. reported LAMB2 to be enriched in the insoluble fraction of the lateral V-SVZ matrisome, suggesting that LAMB2 might be one of the molecules underlying the increased stiffness of the lateral V-SVZ, as compared to other neurogenic and non-neurogenic areas. Given the enrichment of *Lamb2* in slowly versus rapidly dividing RGCs, it would be interesting to assess whether tissue stiffness is increased in the pericellular area of slowly dividing RGCs and whether this could play a role in the emergence of aNSCs. Interestingly, the lateral and more neurogenic V-SVZ is stiffer than the medial and more gliogenic V-SVZ (Kjell et al., 2020). Differences in tissue stiffness might therefore underlie the shift in gliogenic and neurogenic output observed upon *Lamb2* KD, potentially rendering the lateral V-SVZ more similar to the medial V-SVZ in terms of gliogenic/neurogenic potential. Laminins play also a role in anchoring NSCs to their niche by interaction with membrane proteins, such as integrins (Loulier et al., 2009). In this regard, LAMB2 might promote the emergence of aNSCs by stabilizing the attachment of slowly dividing RGCs to the apical VZ surface – where LAMB2 is enriched and where the adult neurogenic niche will be formed – while the MMP15-mediated degradation of LAMB2, and potentially other ECM and cell adhesion molecules, might provide the necessary motility to rapidly dividing RGCs to delaminate and progress toward the neurogenic lineage. Along the same line, the increase of BrdU-retaining NSC-like cells and reduction in neuroblasts following *Mmp15* KD could be caused by impaired delamination of rapidly dividing RGCs, which would force them to enter quiescence. Consistent with this hypothesis, our scRNA-seq highlighted increased percentage of NSC populations characterized by high level of the quiescence marker *Vcam1* and low level of the NSC activation marker *Egfr,* thereby pointing toward increased NSC quiescence upon *Mmp15* KD.

Our study provides novel mechanistic insights into the emergence of aNSCs during development, addressing a largely unexplored function of the extracellular environment in this process and revealing that cells committed to distinct fates – neurogenic NSPCs and future aNSCs – can coexist within the same niche through opposing ECM remodelling strategies.

## Limitation of the study

Although the H3.1-iCOUNT;R26Cre-ERT2 mouse line enables temporal control over the tracing of low- and high-proliferative cells, our single time-point scRNA-seq analysis provides a snapshot of LGE NSPCs rather than a dynamic view of their transcriptional progression. Given that cell cycle dynamics change during LGE lineage progression, we cannot exclude the possibility that some slowly dividing NSPCs identified here will transition to a rapidly dividing state in subsequent cell divisions.

## Data availability

The scRNA-seq raw data generated for this manuscript will be made available on GEO upon publication.

## Acknowledgments

We thank Ines Muehlhahn, Martina Bürkle, Detlef Franzen and Daniel Gonzalez Bohorquez for excellent technical help. We acknowledge the Sequencing Core facility of the Helmholtz Center Munich, the Flow Cytometry Facilities of the Biomedical Center Munich (BMC) and of University of Zurich (UZH) for their services. We are also very grateful to Anthodesmi Krontira, Giacomo Masserdotti, Jovica Ninkovic, Stefan Stricker, Silvia Cappello and Christian Mayer for their valuable input throughout all stages of the project. This work was supported by the European Research Council (advanced grant NeuroCentro, 885382 to M.G) and the German Research Foundation by the Excellence Cluster SyNergy (EXC2145/Project-ID 390857198, to M.G.) and the TRR274 (Nr. 408885537). Parts of this work were funded by a New Frontiers in Research Fund Transformation to M.G., funded through 3 Canadian Federal Agencies (CIHR, NSERC and SSHRC).

## Author contributions

M.G. and D.C. conceived the project and designed the experiments. S.J. and A.D.L. advised on the use of and provided the iCOUNT mouse line and colony for the first experiments. D.C. and A.D.L. performed VASA-seq experiments. A.D. aligned VASA-seq reads, performed RNA velocity, Palantir-based pseudotime and CellRank analysis. D.C. performed all remaining analysis. D.C. performed *in vivo* and *in vitro* experiments. D.C. and P.C. performed immunohistochemistry and western blot. M.T. performed RNA scope. D.C. imaged and quantified the data. D.C. and T.S. prepared 10x library, M.R. performed the alignment and D.C. analyzed the data. D.C. and M.G. wrote the manuscript. M.G. & S.J. acquired funding.

## Declaration of generative AI and AI-assisted technologies in the manuscript preparation process

During the preparation of this work, the author(s) used Claude (Anthropic) to assist with coding and grammar correction of some paragraphs. After using this tool/service, the author(s) reviewed and edited the content as needed and take(s) full responsibility for the content of the published article.

## Materials and Methods

### Mouse husbandry

C57BL/6J mice were bred in the Core Facility Animal Models (CAM) of the BioMedical Center (Ludwig-Maximilians-Universität, Planegg-Martinsried). H3.1-iCount;Rosa26Cre-ERT2 mice were bred in the Animal Facility of the Institute for Brain Research (University of Zurich). The H3.1-iCount mouse strain is available at the Jackson Laboratory (Strain No. 037318). The date of the vaginal plug was considered embryonic day (E) 0 and the date of birth as postnatal day (P) 0. All animal experiments were conducted in accordance with the German animal welfare laws (TierSchG) and guidelines of the GV-SOLAS (Society of Laboratory Animal Science) – for C57BL/6J mice – or in accordance with Swiss regulatory standards and approved by the Veterinary office of the Canton of Zurich – for H3.1-iCount;Rosa26Cre-ERT2 mice. Mice were kept in individually ventilated cages under a 12-hour light/dark cycle with *ad libitum* access to water and food.

### Fluorescence-activated cell sorting (FACS) of embryonic tissue for VASA-seq

Double homozygous pregnant H3.1-iCount;Rosa26Cre-ERT2 females received an intraperitoneal injection of tamoxifen (180 mg/kg, Sigma-Aldrich) dissolved in corn oil (Sigma-Aldrich) at E14.5, and embryonic lateral ganglionic eminences (LGEs) were collected two days post-injection (E16.5). For plates 1 and 2, LGEs were manually dissected from five embryos (one litter) in ice-cold 1x HBSS (Thermo Fisher Scientific) supplemented with 10 mM HEPES, centrifuged at 1,000 rpm at 4°C for 5 minutes, and enzymatically dissociated in 1 mL of 0.05% Trypsin/EDTA for 15 minutes at 37°C. Enzymatic activity was inhibited with 1**×** HBSS supplemented with 10% fetal bovine serum (FBS), followed by mechanical dissociation using a fire-polished glass Pasteur pipette. For plates 3 and 4, LGEs were manually dissected from eight embryos (one litter) under the same conditions and dissociated using the Neural Tissue Dissociation Kit (Miltenyi Biotec, Cat. No. 130-092-628) according to the manufacturer’s instructions. For plate 5, LGEs were dissected from six embryos (two litters) in ice-cold 1**×** HBSS, embedded in 4% low-melting point agarose, and sectioned at 200-300 μm using a vibratome. LGE germinal zones were subsequently microdissected and processed as described for plates 1 and 2. Following dissociation, cell pellets were resuspended in 1**×** DPBS (Thermo Fisher Scientific) or 1**×** HBSS (Thermo Fisher Scientific) containing RNase inhibitors (RNasin; Promega, Cat. No. N2511) and filtered through a 40 μm cell strainer (Sigma-Aldrich). Cell viability was assessed using Zombie Violet (BioLegend, Cat. No. 423113) at a 1:1,000 dilution. Fluorescence-activated cell sorting was performed on a FACSAria™ III Cell Sorter (BD Biosciences). GFP− and mCherry-expressing cells were identified using the FITC-A and PE-Texas Red-A channels, respectively, with brain tissue from wild-type C57BL/6 mice used as negative control for FACS gating. GFP single-positive and mCherry/GFP double-positive cells were sorted into 384-well cell capture plates, pre-filled with CEL-seq2 barcodes and primers, supplied by Single Cell Discoveries (https://www.scdiscoveries.com/services/vasa-seq/). Plates were briefly centrifuged, snap-frozen on dry ice, and stored at −80°C.

### VASA-seq library preparation

Library preparation and single-cell RNA sequencing from H3.1-iCount;Rosa26Cre-ERT2 samples were carried out by Single Cell Discoveries using the VASA-seq technology (Salmen et *al.*, 2022). Briefly, following single-cell lysis, total RNA was fragmented and the resulting fragments were end-repaired and polyadenylated. Polyadenylated fragments were then reverse-transcribed into cDNA using barcoded oligo-dT primers, rendered double-stranded, and amplified by in vitro transcription. Ribosomal RNA was depleted from the amplified RNA (aRNA) pool, and sequencing libraries were finalized through ligation, reverse transcription, and PCR amplification, yielding UMI-containing fragments that were subsequently sequenced on an Illumina NextSeq 500 platform at a sequencing depth of 150,000 reads per cell.

### Fluorescence-activated cell sorting (FACS) of postnatal tissue for 10x sequencing

Brain tissues from P10 pups, injected at E13 with lentiviral vectors carrying microRNA versus *Mmp15* or a non-targeting microRNA, were collected in ice-cold 1**×** HBSS and immediately sectioned using a brain matrix in order to obtain 500 μm coronal sections. Next, sections were placed in ice-cold dissection buffer (1**×** HBSS supplemented with 10 mM HEPES) and the lateral V-SVZ was manually dissected and collected in dissection buffer. Samples were centrifuged at 1,000 rpm at 4°C for 5 minutes and enzymatically dissociated in 1 mL of 0.05% Trypsin/EDTA for 15 minutes at 37°C. Enzymatic activity was inhibited with 1**×** HBSS supplemented with 10% fetal bovine serum (FBS), followed by mechanical dissociation using a fire-polished glass Pasteur pipette. Single-cell suspension was centrifuged at 1,000 rpm at 4°C for 5 minutes and resuspended in 1**×** PBS containing Zombie Violet (BioLegend, Cat. No. 423113) at a 1:1,000 dilution, for 15 minutes at 4°C. Next, cells were washed in 1**×** PBS1, resuspended in 1**×** HBSS supplemented with 1 % bovine serum albumin (BSA, Miltenyi Biotech, Cat. No 130-091-376) and filtered through a 40 μm cell strainer. Brain tissue from wild-type C57BL/6 pups was used as negative controls for FACS gating. FACS was performed at 4°C on a FACSAria™ III Cell Sorter (BD Biosciences). GFP+ and GFP− cells were collected into 1.5 mL protein low-binding tubes pre-coated with sorting buffer (1 % BSA in 1**×** HBSS) for 3 hours at 37°C and pre-filled with 100 μl of sorting buffer. Samples were then centrifuged at 300 g at 4°C for 10 minutes and resuspended in 30-20 μl of sorting buffer. GFP+ and GFP− cells were mixed in a 1:2 proportion and processed for 10x library preparation.

### 10x library preparation

10x library preparation was performed using the Chromium GEM-X Single Cell 3’ v4 Gene Expression Kit (PN-1000686), according to manufacturer’s instructions. Briefly, single-cell suspensions were loaded onto a GEM-X Chip v4 (PN-1000690) to partition cells into nanoliter-scale Gel Bead-in-Emulsions (GEMs), where cell barcoding and reverse transcription occurred simultaneously. Next, barcoded cDNA was purified, amplified by PCR, and processed into 3’ gene expression libraries through enzymatic fragmentation, end repair, adaptor ligation, and dual-index PCR (Dual Index Kit TT Set A, PN-1000215). Library quality and quantity were assessed prior to sequencing. Samples were sequenced on an Illumina NovaSeq X Plus platform, at a sequencing depth of ∼80.000 reads per cell, at the Genomics Core Facility of the Helmholtz Center Munich.

### Transcriptome analysis

VASA-seq reads were aligned against the mouse genome (GRCm39, UCSC Genome Browser, version M35), with the GFP and mCherry sequences included, using *zUMIs* pipeline (Parekh et al., 2018). The resulting count matrices were analysed using Seurat v5 standard workflow. Following quality control, cells with a total number of detected genes (nFeature_RNA) falling outside the 12th and 98th percentiles or nCount_RNA below the 12th percentile of the sample distribution were discarded. Normalization was performed using the function “NormalizeData” and cell-cycle scores were assigned via the Seurat function “CellCycleScoring”, using a list of cell-cycle related genes (Tirosh et al., 2016). Differences in G2M- and S-phase scores were regressed out during genes scaling using the function “vars.to.regress = (’CC.Difference’)”. Datasets from different 384-well plates were integrated using CCA integration methods. Clusters were identified using the “FindClusters” function (dimensions = 20, res = 1) and manually annotated based on the expression of known marker genes. RNA velocity was performed on scVelo (Bergen et al., 2020) using the dynamical model computed together for rapidly and slowly dividing cell populations. Pseudotime analysis was computed independently for slowly and rapidly dividing cells using Palantir 1.4.4 (Setty et al., 2019). Fate probability analysis was performed using CellRank 2.1.0. (Lange et al., 2022), setting RGCs as initial state and neuronal clusters as terminal states. Differential expression (DE) analysis between slowly and rapidly dividing NSPCs was performed using the function “FindMarkers” with default parameters setting ident.1 = “Slowly dividing cells” and ident.2 = “Rapidly dividing cells”, and min.pct = 0.3. Cells in cluster IPs_3 were downsampled to obtain equal numbers of slowly and rapidly dividing cells for DE analysis. Differentially expressed genes (DEGs) were visualized using EnhancedVolcano R package (Blighe et al., 2025), using *p-value* < 0.05 and avg_log2FC > 0.5 as thresholds. Gene ontology (GO) analysis was performed using clusterProfiler (Yu et al., 2012) with the following settings: ont = “BP”, pAdjustMethod = “BH”, pvalueCutoff = 0.05, qvalueCutoff = 0.1. All the genes detected in our dataset were used as background genes for GO analysis. GO terms shown in Fig. 2 and Fig. S2 were manually selected among the enriched biological process with qvalue < 0.1 for RGCs and IPs_1, qvalue < 0.3 for IPs_2 and IPs_3. GO term analysis on the shared enriched biological processes among slowly dividing RGCs and pNSCs was performed as described above, using as background all the genes detected in our and Cebrian-Silla et al., 2021 datasets.

10x Genomics reads were aligned against the mouse genome (GRCm39, UCSC Genome Browser), with the GFP and Woodchuck Hepatitis Virus Posttranscriptional Regulatory Element (WPRE) sequences included, using CellRanger v9.0.1. The resulting count matrices were analysed using Seurat v5 standard workflow. Control and *Mmp15* KD count matrices were merged and subjected to quality control. Cells with less than 10% mitochondrial genes, more than 1.000 and less than 10.000 number of detected genes (nFeature_RNA) were retained for downstream analysis. Blood cells were manually removed and doublets were computationally identified and removed using the package DoubletFinder (McGinnis et al., 2019). Normalization and cell cycle regression were performed as described above. Clusters were identified using the “FindClusters” function (dimensions = 12, res = 1) and manually annotated based on the expression of known marker genes. Infected and not-infected cells were identified based on the expression of the WPRE sequence on the unintegrated viral genome, using the function “WhichCells” and setting “expression = WPRE > 0” or “expression = WPRE == 0”, respectively.

### Brain tissue preparation

For fluorescence in situ hybridization via RNA scope, embryonic brains were dissected in ice-cold 1× PBS and fixed in 4% PFA (Carl Roth) at 4°C for durations adjusted to the developmental stage (1 hour for E12, 2 hours for E14, 4 hours for E16, and 6 hours for E18). Following fixation, tissues were rinsed in 1× PBS and subsequently cryoprotected by overnight immersion in 30% sucrose in 1× PBS at 4°C. Brains were then embedded in Neg-50 Frozen Section Medium (Epredia™, Thermo Fisher Scientific, Cat. No. 6502), rapidly frozen on dry ice, and sectioned coronally at 25-30 μm thickness on a cryostat.

For immunohistochemistry, pups were anesthetized via injection of Medetomidine (0.5 mg/kg), Midazolam (5 mg/kg), and Fentanyl (0.05 mg/kg) and transcardially perfused with ice-cold 1× PBS followed by ice-cold 4% PFA. Postnatal brains were dissected and post-fixed in 4% PFA at 4°C overnight. 50 μm coronal sections were obtained using a vibrotome. Both male and female pups were used for immunohistochemistry.

### RNA scope

Expression of *Lamb2* and *Mmp15* during brain development was assessed by fluorescence in situ hybridization using the RNAscope Multiplex Fluorescent Detection Kit v2 (ACD, Cat. No. 323110), together with the following associated reagents: Pretreatment Reagents (ACD, Cat. No. 322381), Multiplex TSA Buffer (ACD, Cat. No. 322810), Target Retrieval Reagents (ACD, Cat. No. 322000), and Wash Buffer Reagents (ACD, Cat. No. 310091). All procedures were carried out following the manufacturer’s protocol with minor modifications. Briefly, cryosections were rinsed in 1× PBS for 5 minutes at room temperature (RT), dried at 60°C for 30 minutes, and post-fixed in 4% PFA (Carl Roth) for 15 minutes at 4°C. Tissue dehydration was achieved by sequential 5-minute immersions in 50%, 70%, and twice in 100% ethanol (Merck Millipore) at RT. Endogenous peroxidase activity was quenched by treatment with hydrogen peroxide for 10 minutes at RT, followed by rinsing with DEPC-treated water (diethyl pyrocarbonate, SERVA). Antigen retrieval was performed by immersing slides in boiling retrieval solution (99°C) for 3 minutes, after which sections were rinsed with DEPC-water and briefly dehydrated in 100% ethanol. Proteolytic digestion was carried out using Protease III for 10 minutes at 40°C, followed by a brief DEPC-water rinse. Target probes – RNAscope™ Probe-Mm-Lamb2-C2 (ACD, Cat. No. 412891-C2) and RNAscope™ Probe-Mm-Mmp15-C3 (ACD, Cat. No. 1283231-C3) – were diluted 1:50 in probe dilution reagent (ACD, Cat. No. 300041) and hybridized for 2 hours at 40°C. A 3-plex negative control probe (ACD, Cat. No. 320871) and a 3-plex positive control probe (ACD, Cat. No. 320881) were included in every experimental run. After hybridization, slides were washed twice in 1× wash buffer for 2 minutes at RT, and signals were amplified through successive incubations with AMP 1 (30 minutes, 40°C), AMP 2 (30 minutes, 40°C), and AMP 3 (15 minutes, 40°C). Fluorescent detection was achieved by incubating sections with HRP-C2 (for *Lamb2*) or HRP-C3 (for *Mmp15*) for 15 minutes at 40°C, followed by a 30-minute incubation at 40°C with Opal-570 fluorophore (1:750; Akoya Biosciences, Cat. No. FP1488001KT) diluted in Multiplex TSA Buffer. HRP activity was subsequently blocked by incubation with HRP Blocker for 15 minutes at 40°C, and slides were washed twice in 1× wash buffer. Nuclei were counterstained with DAPI (1:1000; Sigma-Aldrich).

### Recombinant DNA cloning

MicroRNAs sequences were designed using the Block-iT™ RNAi Designer tool by Invitrogen (rnaidesigner.thermofisher.com/rnaiexpress) and cloned in the plasmid pENTRY-GW-grandestuffer_GFP, containing the cDNA of the Emerald green fluorescent protein (EmGFP). The resulting plasmids were subcloned into retroviral or lentiviral transfer plasmids under a CAG promoter using the Gateway clonase system (Invitrogen, Cat. No. 11791020), according to the manufacturer’s protocol. The following microRNA sequences were used: 5’-TGCTGAAATGTACTGCGCGTGGAGACGTTTTGGCCACTGACTGACGTCTCCACGCAGTACATTTC-3’(non-targeting), 5’-TGCTGTTATTGAAAGCTCTTGCTAGCGTTTTGGCCACTGACTGACGCTAGCAAGCTTTCAATAA-3’(Lamb2), 5’-TGCTGTGAGCATCCACAATCACACACGTTTTGGCCACTGACTGACGTGTGTGAGTGGATGCTCA-3’ (Mmp15).

### Viral production

For lentiviral production, pVSVG envelope plasmid, pCMV-dR8.91 packaging helper plasmid and lentiviral transfer plasmids (carrying GFP reporter and microRNA sequences), were co-transfected into HEK283T cells. For retroviral production, retroviral plasmids (carrying GFP reporter and microRNA sequences) were transfected into GPG293 retroviral packaging cell line. Supernatant containing viral particles was collected 48-72 hours post-transfection. The viral titer was calculated as transduced units per ml (TU/ml), via infection of postnatal mouse astrocytes (retroviral vectores) or HEK283T cells (lentiviral vectors) with serial dilutions of viral suspension. Viral stocks were used at concentrations ranging from 10¹¹ to 10¹³ TU/mL, with comparable amounts applied to both control and knockdown conditions.

### Cell transfection

Neuro2a (N2A) cells (ATCC, Cat. No. CCL-131) were maintained on uncoated plates in growth medium consisting of DMEM + GlutaMax supplemented with 10% fetal bovine serum (FBS), 1% penicillin/streptomycin, 1% sodium pyruvate, and 1% non-essential amino acids (NEAA), at 37°C in a humidified atmosphere containing 5% CO₂. Upon reaching confluency, cells were passaged by rinsing with 1× PBS followed by mechanical dissociation. To assess the knockdown efficiency of miRNAs targeting *Lamb2* and *Mmp15*, N2A cells were transfected with viral transfer plasmids using Lipofectamine™ 2000 Transfection Reagent (Thermo Fisher Scientific, Cat. No. 11668027) following the manufacturer’s instructions. Untreated cells and mock-transfected cells (i.e. cells exposed to transfection reagent in the absence of plasmid DNA) were included as experimental controls. Three days after transfection, cells were mechanically dissociated, transfection efficiency was measured via FACS based on the expression of GFP reporter and cells lysed for protein collection.

### Protein extraction and quantification

N2A cells were lysated in RIPA buffer (Sigma-Aldrich Cat. No. R0278) containing 1× protease inhibitor (cOmplete Protease Inhibitor Cocktail, Roche Cat. No. 11697498001). Samples were then incubated 15 minutes in ice and centrifuged at 13.000 g at 4°C for 15 minutes. The supernatant was collected and protein concentration was determined via Bradford assay.

### Degradation assay

Recombinant Human MMP-15/MT2-MMP CF (R&D Systems, Cat. No. 916-M) was activated according to manufacturer’s instructions with minor modifications. Briefly, 1 μg of recombinant MMP15 protein was incubated with 0.1 μg/mL Trypsin in activation buffer (50 mM Tris, 10 mM CaCl2, 150 mM NaCl, 5 µM ZnCl2, 0.05% (w/v) Brij-35, pH 7.5) for 1 hour at 37°C. Trypsin-mediated digestion was inhibited by adding 4-(2-Aminoethyl) benzolsulfonylfluorid (AEBSF, Tocris, Cat. No. 5175) at a final concentration of 1mM for 15 minutes at RT. Next, activated MMP-15 was diluted to 10 ng/µL in Assay Buffer (50 mM Tris, 500 mM NaCl, 5 mM CaCl_2_, 1 µM ZnCl_2_, 0.02% (w/v) Brij-35, pH 8.0). 0, 30 or 60 ng of activated MMP15 were incubated with 500 ng of full-length human recombinant laminin-521 (Biolamina, Cat. No. LN-521) 4 hours at 37°C. Samples were then stored at −20°C.

### Western blot

Protein samples were mixed with 1× Laemmli buffer and denatured by boiling at 95°C for 5 minutes. Proteins of different molecular weights were separated by SDS-PAGE on 8% polyacrylamide gels and subsequently transferred to a nitrocellulose membrane. The membrane was then blocked with 5% non-fat dry milk in 1× TBS-T (0.1% Tween-20 in Tris-buffered saline, pH 7.4) for 1 hour at RT to prevent non-specific antibody binding. Primary antibodies were diluted in blocking solution and incubated with the membrane overnight at 4°C. The following day, membranes were washed and incubated with HRP-conjugated secondary antibodies diluted in blocking solution for 1 hour at RT. Signal was detected using enhanced chemiluminescence (ECL). The following primary antibody were used: anti-LAMB2 (Proteintech Cat. No. 30943-1-AP) 1:1000 and 1:2000, anti-MMP15 (MyBioSource, Cat. No. MBS1753732) 1:1000, anti-GAPDH (Santa Cruz, Cat. No. sc32233) 1:10.000.

### In utero injection (IUI)

All animal experiments were conducted under license by the Government of Upper Bavaria. E13 timed-pregnant C57BL/6J mice were anesthetized via intraperitoneal injection of Medetomidine (0.5 mg/kg), Midazolam (5 mg/kg), and Fentanyl (0.05 mg/kg). 1 μL of titrated viral solution containing 0.01% Fast Green dye was injected into the lateral ventricle of each embryo using a glass capillary of approximately 10 µm diameter (Drummond Scientific Cat. No. 5-000-1001-X10). The uterus was carefully repositioned within the abdominal cavity which was subsequently closed with surgical sutures. Anesthesia was reversed by subcutaneous administration of Buprenorphine (0.1 mg/kg), Atipamezole (2.5 mg/kg), and Flumazenil (0.5 mg/kg).

### BrdU administration

For detection of slowly dividing cells in the postnatal ventricular-subventricular zone (V-SVZ) of C57BL/6J mice, 5-bromodeoxyuridine (BrdU; Sigma-Aldrich, Cat. No. B5002-5G) was dissolved in 0.9% NaCl and administered intraperitoneally (100 mg/kg) to pregnant females at E13.

### Immunostainings

Brain sections rinsed twice in T-PBS containing 1× PBS and 0.25% Triton X-100 (Sigma-Aldrich, Cat. No. T8787) for 10 minutes per wash, followed by a 1-hour incubation in blocking solution, consisting of 3% bovine serum albumin (BSA; Sigma-Aldrich, Cat. No. A2153) and 0.5% Triton X-100 (Sigma-Aldrich, Cat. No. T8787) in 1× PBS, at RT. Primary antibodies diluted in blocking solution were applied overnight at 4°C. Sections were subsequently washed twice in T-PBS (10 minutes per wash) and incubated with secondary antibodies and DAPI (1:1000; Sigma-Aldrich) for 1 hour and 30 minutes at RT. Finally, sections were washed three times in 1× PBS (10 minutes per wash) and coverslipped using Aqua Poly/Mount medium (Polysciences, Cat. No. 18606) on Epredia™ microscope slides. BrdU staining was performed in three sequential days. Briefly, sections were washed twice in T-PBS (10 minutes per wash), incubated in 3N HCl (Carl Roth, Cat. No. 4625.1) for 30 minutes at RT to denature DNA, and HCl neutralized by two washes in 0.1 M boric acid (15 minutes per wash). Sections were then blocked for 1 hour at RT and incubated overnight at 4°C with primary antibodies against BrdU and GFP. The following day, sections were washed twice in T-PBS (10 minutes per wash) and incubated with the corresponding secondary antibodies for 1 hour and 30 minutes at RT. Sections were then washed twice in T-PBS (10 minutes per wash), post-fixed in 4% PFA for 10 minutes at RT, washed twice again in T-PBS (10 minutes per wash), and subsequently incubated with primary and secondary antibodies against GFAP as described above. The following primary antibody were used: anti-GFP (AvesLab, Cat. No. GFP-1020) 1:500, anti-GFAP (Dako, Cat. No. Z0334) 1:250, anti-BrdU (Abcam, Cat. No. AB6326) 1:500, anti-DCX (Santa Cruz, Cat. No. sc-8066) 1:500, anti-DCX (Merck/Millipore, Cat. No. AB2253) 1:1000, anti-OLIG2 (Merck/Millipore, Cat. No. MABN50 and AB9610) 1:250 and 1:500, respectively, anti-KI67 (Abcam, Cat. No. AB15580) 1:500.

### Imaging, quantification and statistical analysis

Brain sections were imaged by confocal laser scanning microscopy (Zeiss LSM710 or Leica Stellaris) using a 40× objective. Images were acquired as z-stacks of 15–16.5 µm depth with a step size of 1–1.5 µm. For each condition, 2–4 coronal sections from 3–9 postnatal brains were analysed, with pups derived from at least 2 independent litters. The V-SVZ was defined as the region of DAPI-dense nuclei lining the lateral ventricles, plus 4-5 nuclei above. The rostral migratory stream (RMS) was excluded from the analysis. GFP+ cells were quantified across the entire z-stack depth. Quantifications were performed using ImageJ (NIH), and statistical analyses were carried out using GraphPad Prism 8.0. Prior to statistical testing, normality of the data distribution was assessed using the Shapiro–Wilk test. Statistical tests are specified in the respective figure legends.

## Supplemental figure legends

**Suppl. Figure 1.**
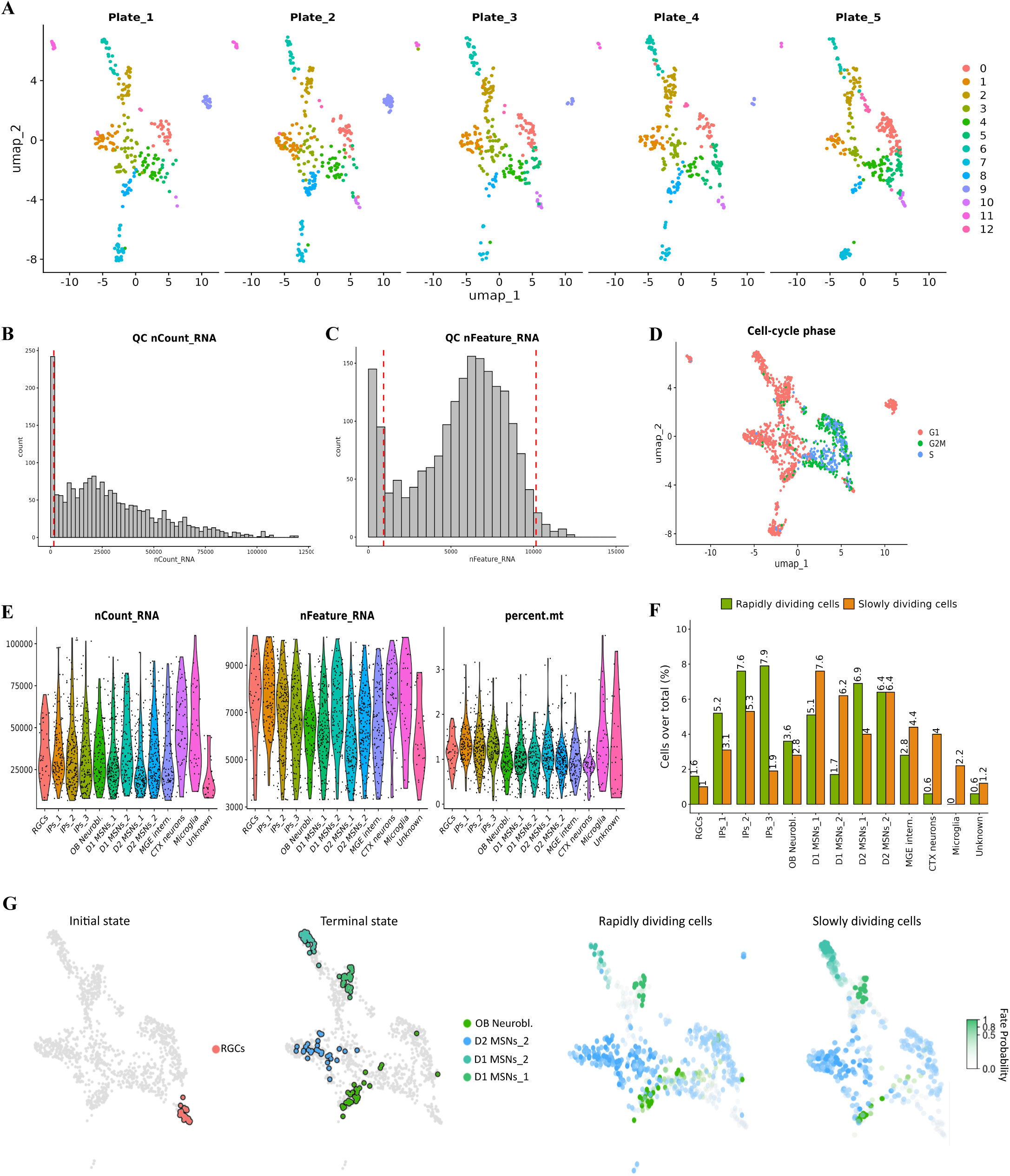
Quality control and transcriptome analysis of LGE cells sorted from iCOUNT mice. A. UMAP plots showing distribution of cell population across different 384-well plates following integration. B-C. Histograms showing distribution of counts and features in integrated datasets and thresholds (red dotted lines) used for the quality control. D. UMAP plot showing cell cycle scores across clusters after cell cycle regression. E. Violin plots showing distribution of counts, features and mitochondrial genes across clusters. Note that cluster ‘unknown’ has lower counts and features and higher mitochondrial gene expression compared to other clusters. F. Histogram showing the proportion of slowly and rapidly dividing cells in each cluster over the total clusters. Note that *Tbr1*-expressing cells mainly derived from low proliferative cells. G. CellRank fate probability analysis showing manually selected initial and terminal states, and fate probabilities for slowly and rapidly dividing cells. Each cell is colored according to its most likely fate; color intensity reflects the degree of lineage priming.

**Suppl. Figure 2.**
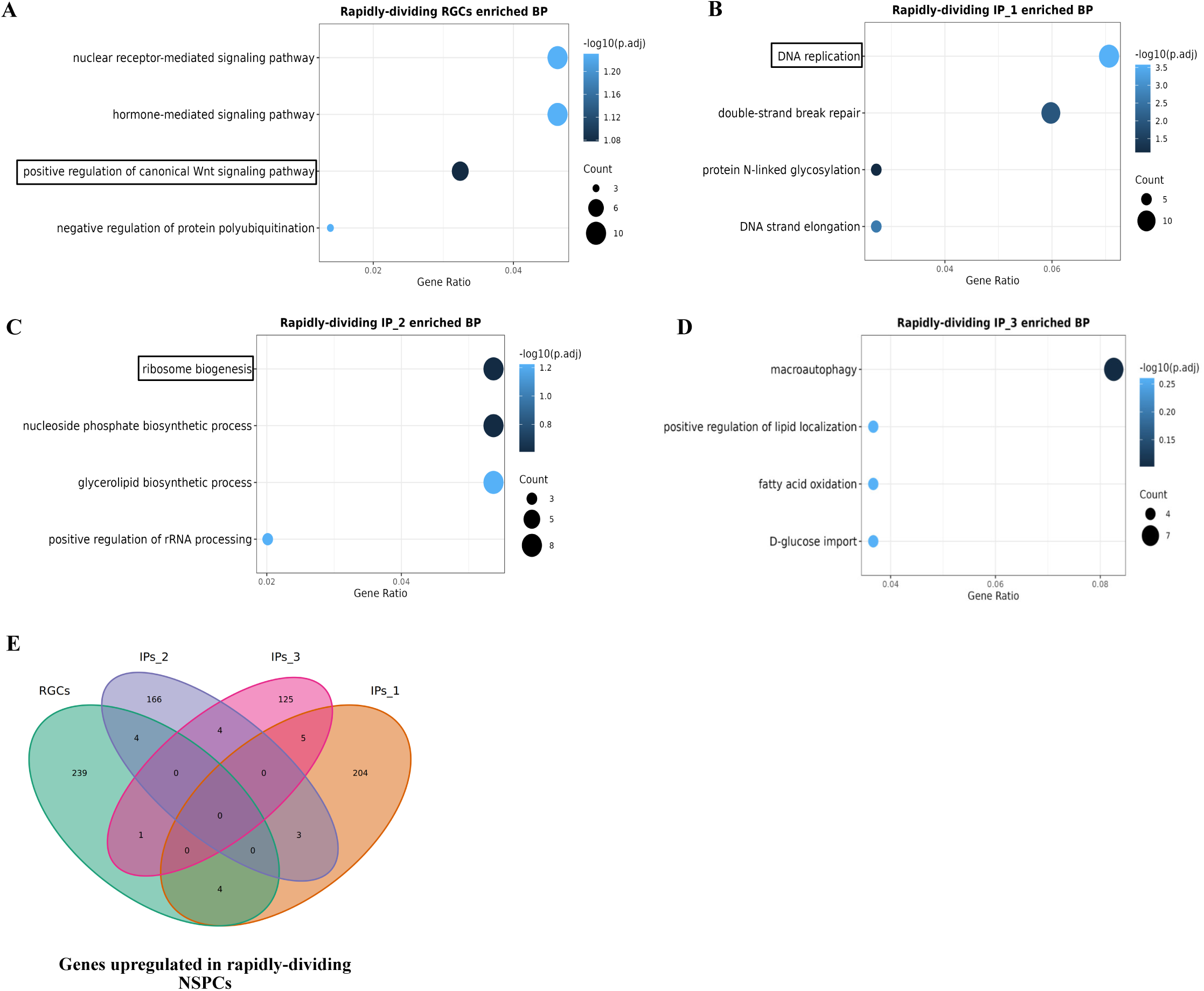
Biological processes enriched in rapidly dividing NSPCs. A-D. Selected biological processes enriched in rapidly dividing RGC and IP clusters. Note that RGCs, IPs_1 and IPs_2 show biological processes – such as Wnt signaling, DNA replication and ribosome biogenesis – linked to cell proliferation. E. Venn diagram showing overlap in rapidly dividing cells-enriched DEGs across RGC and IP clusters.

**Suppl. Figure 3.**
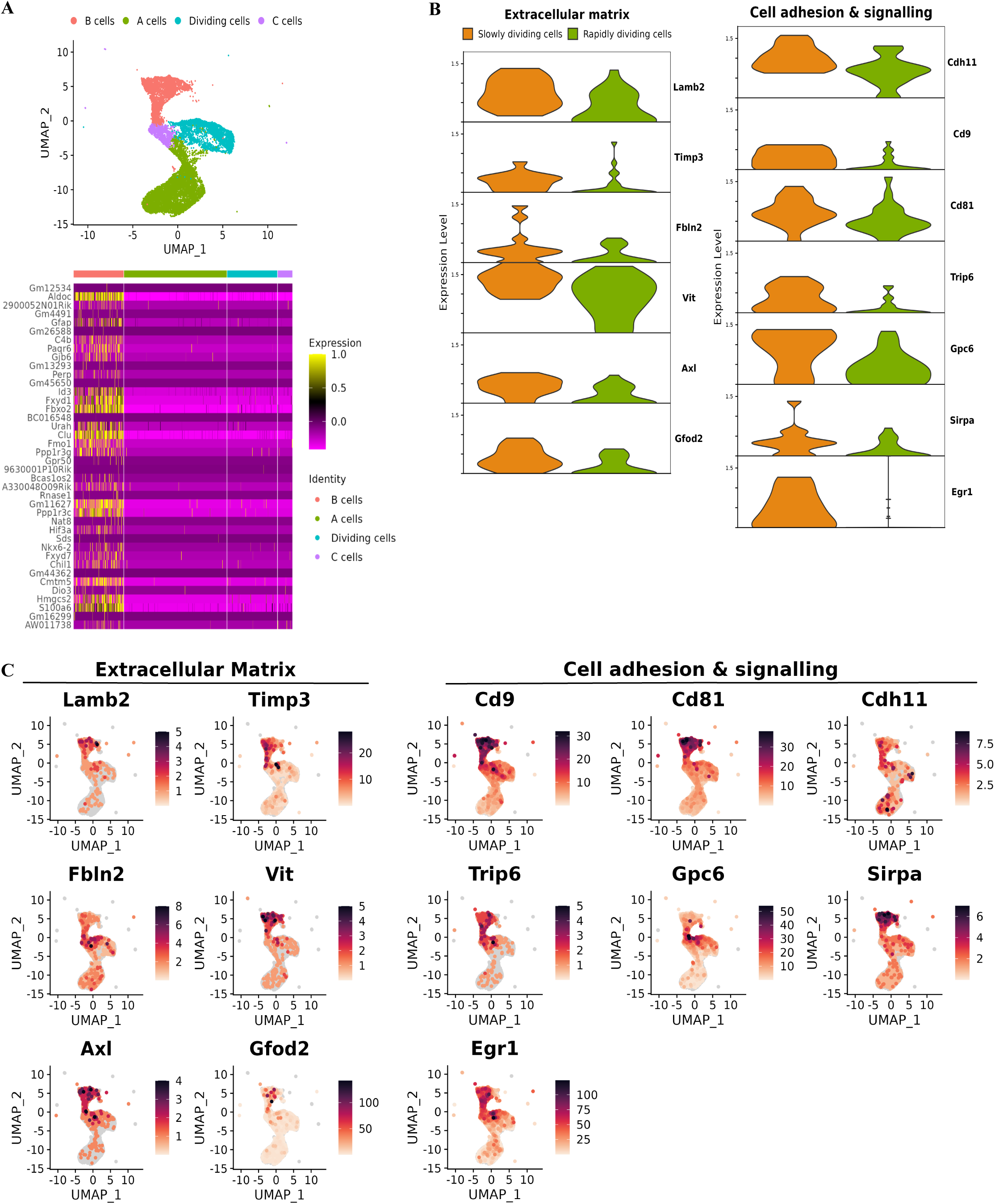
Gene signature of pNSCs and expression of ECM- and cell adhesion-associated genes in embryonic RGCs and postnatal V-SVZ cells. A. UMAP from Cebrian-Silla et *al.*, 2021 showing cell populations contributing to the neurogenic lineage. Postnatal NSCs gene signature was created by performing DE analysis between NSCs (B cells), TAPs (here considered as “A cells” and “dividing cells”) and neuroblasts (C cells), as shown in the heatmap. B. Violin plots showing enrichment of ECM- and cell adhesion-associated genes – shared in slowly dividing RGCs and pNSCs – in slowly versus rapidly dividing RGCs. C. UMAP plots showing enrichment of ECM- and cell adhesion-associated genes – shared in slowly dividing RGCs and pNSCs – in pNSCs as compared to the other cell clusters.

**Suppl. Figure 4.**
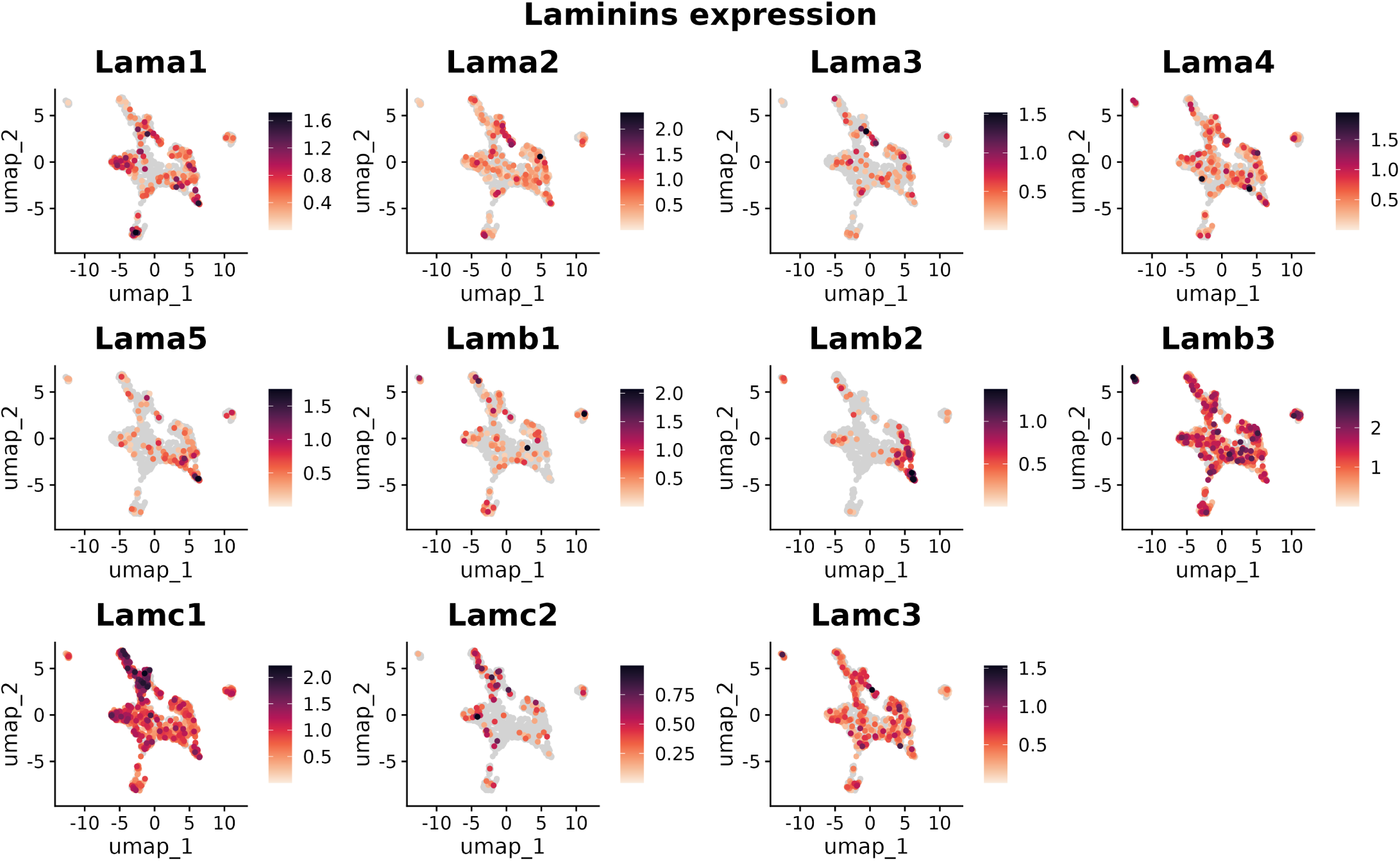
Expression of laminin chain isoforms across sorted LGE cell population from iCOUNT mice. UMAP showing normalized expression of all known laminin chain isoforms across clusters. Note that only *Lamb2* and *Lama5* show enriched expression in the RGC population as compared to other clusters.

**Suppl. Figure 5.**
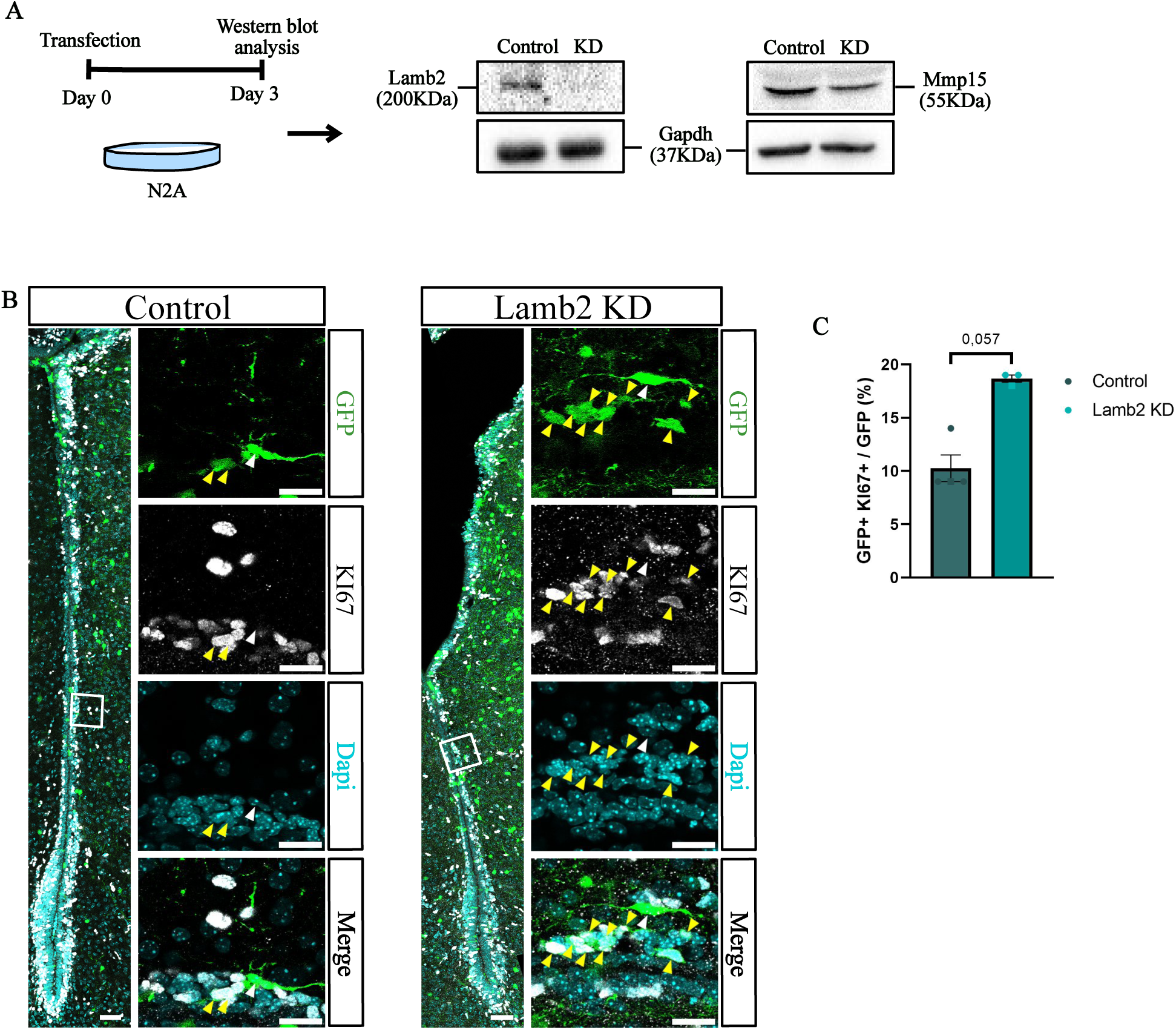
*Lamb2* and *Mmp15* KDs efficiency and effect on cell proliferation following *Lamb2* KD. A. *Lamb2* and *Mmp15* KDs efficiency was tested in vitro by transfection of N2A cells with plasmids carrying microRNAs versus the target genes or a control non-targeting microRNA Three days after transfection proteins were extracted and KD efficiency was assessed by western blot. B-C. Representative images and quantification of GFP+ KI67+ cells in the P10 lateral V-SVZ following *Lamb2* KD. Yellow and white arrows indicate cells positive or negative for KI67, respectively. Scale bar: 50μm in the overview images, 20μm in high-magnification images. Data are shown as means ± SEM. Dots with different shapes correspond to pups from different litters. Mann-Whitney Test.

**Suppl. Figure 6.**
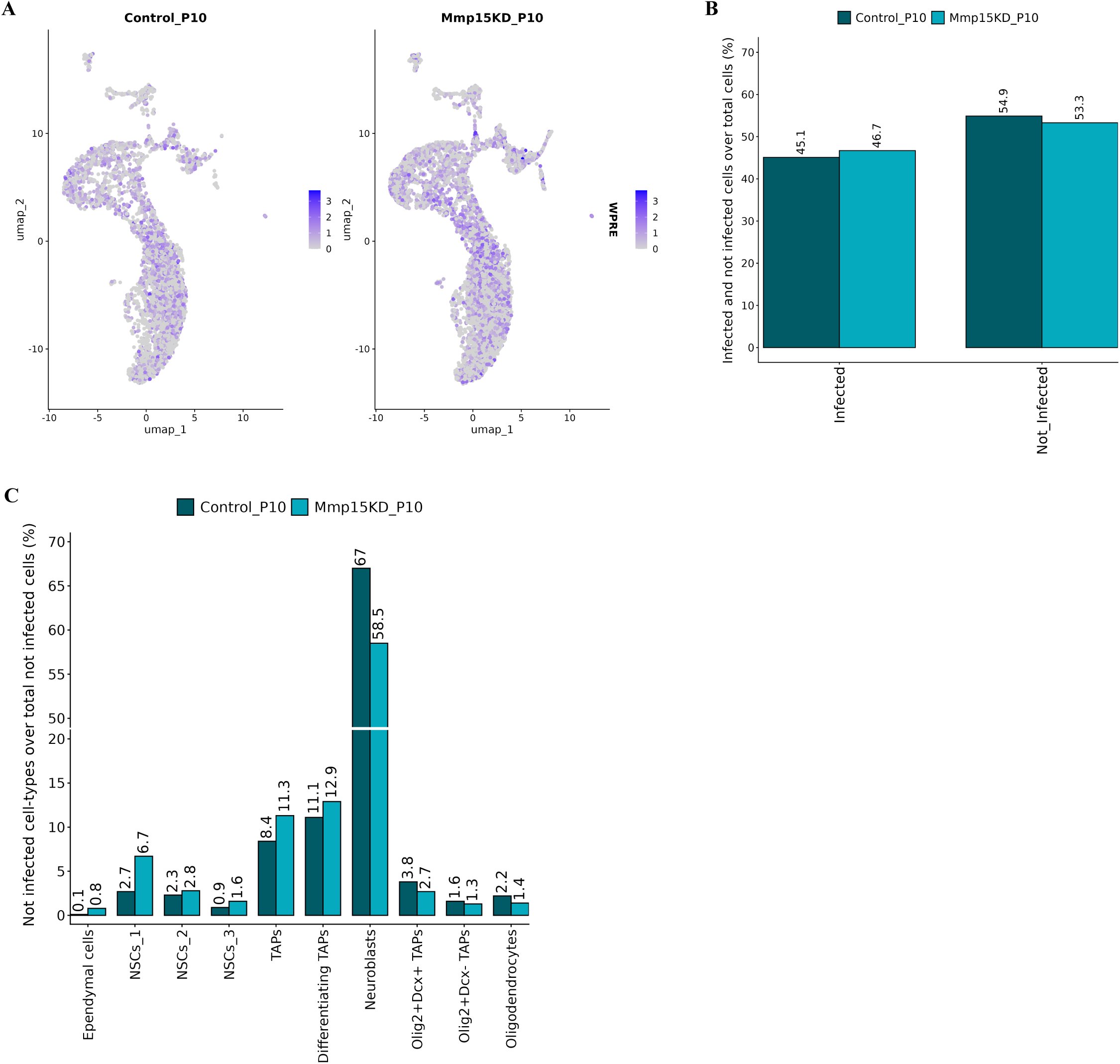
Identification of infected and not infected cells in scRNA-seq of P10 V-SVZ cells following *Mmp15* KD. A. Feature Plot showing expression of WPRE in control and *Mmp15* KD datasets. B. Histogram showing the percentage of both infected and not-infected cells in control and *Mmp15* KD datasets. Note that proportions are similar across conditions. C. Histogram showing the percentage of not-infected cell-types over all the non-infected cells.

